# Nanobodies targeting hnRNPA2/B1 and tau

**DOI:** 10.1101/2025.11.21.689763

**Authors:** Azady Pirhanov, Cristian Rodriguez, Fatemeh Tashakori-Asfestani, Raven Gonsoulin, Rayne Santiago, Katarnut Tobunluepop, Emmanuella Erhunmwunsee, Sambhavi Puri, Monika Arbaciauskaite, Wu He, Benjamin Wolozin, Yongku Cho

## Abstract

Heterogeneous nuclear ribonucleoprotein A2/B1 (hnRNPA2/B1) is an RNA-binding protein that mislocalizes to the cytoplasm and forms stress-induced granules in tauopathy and multisystem proteinopathy. It also preferentially interacts with oligomeric tau and is required for tau-mediated neurodegeneration in a mouse model of Alzheimer’s disease. To study endogenous hnRNPA2/B1 and tau, we generated nanobodies that specifically recognize these proteins. We screened yeast surface display nanobody libraries using an avidity-enhanced screening strategy that enabled selection of binders against short peptide ligands. This led to isolation of anti-hnRNPA2/B1 and anti-tau nanobodies with defined epitopes. Directed evolution of the anti-hnRNPA2/B1 nanobody improved binding affinity by over 20-fold but caused cytoplasmic aggregation, demonstrating a tradeoff between affinity and intracellular behavior. Although the final nanobodies retained modest affinities, they showed robust intracellular colocalization with their targets. Furthermore, fusion to ubiquitin ligase adaptor domains significantly decreased hnRNPA2/B1 and tau levels. Collectively, these nanobodies provide valuable tools for studying hnRNPA2/B1 and tau dynamics in their native cellular context.

## 1. Introduction

RNA-binding proteins (**RBPs**) have emerged as key players in the pathology of neurodegenerative diseases, including Alzheimer’s disease (**AD**), amyotrophic lateral sclerosis (**ALS**), and frontotemporal dementia (1–3). Heterogeneous nuclear ribonucleoprotein A2/B1 (**hnRNPA2/B1**) is an RBP involved in pre-mRNA processing, splicing, mRNA trafficking, and post-transcriptional gene expression control (4). A mutation in hnRNPA2/B1 (D290V) has been discovered in a family with multisystem proteinopathy, a degenerative disease affecting muscle, bone, and the central nervous system (5, 6). We recently showed that hnRNPA2/B1 interacts with oligomerized tau and is required for tau-mediated neurodegeneration in a mouse model of AD (7).

Many studies have reported changes in RBP localization and dynamics in the diseased state (8–11). In particular, RBPs that have a low complexity domain have a propensity to form phase-separated condensates (12, 13). Disease-linked mutations in RBPs alter the phase-separation properties of the condensates, typically making them more amyloidogenic and reducing their dynamics (14–16). hnRNPA2/B1 is mislocalized to the cytoplasm in multi-system proteinopathy, ALS, and tauopathies (7, 17–19). hnRNPA2/B1 undergoes phase separation *in vitro* (20, 21), but whether this phenomenon plays a role in neurodegeneration is yet to be determined.

These findings demonstrate the importance of studying the dynamics of these proteins in vivo and yet also illustrate the challenges in investigating the dynamics of endogenous cellular constituents. Since condensate formation is driven by an increase in concentration (22), current methods that rely on the expression of fluorescently tagged components may promote phase separation. Therefore, monitoring the constituents without perturbing the endogenous concentration is highly advantageous in studying RBPs.

Nanobodies (**Nbs**) have a strong potential to address this need. Nbs are the variable region of heavy-chain only antibodies from camelids (23, 24). Owing to their small size, favorable stability, and high paratope diversity (25), Nbs have been widely used in many intracellular targeting applications, including structural biology (26), virus neutralization (27), targeted protein degradation (28), G protein-coupled receptor modulation (29, 30), kinase regulation (31), and live cell/tissue imaging (32, 33). Using Nbs that bind to RBPs will allow imaging RBP dynamics without increasing RBP concentration. Moreover, by using Nb-targeted degradation strategies, RBP concentrations can be selectively lowered, which may prevent or reverse condensate formation. However, the dominantly unstructured nature and prevalent homology between RBPs make identifying RBP-specific Nbs challenging.

Here, we report a multivalent Nb screening approach that enabled the screening of Nbs against the RBP hnRNPA2/B1 and the tau protein. Taking advantage of the high avidity of yeast surface display (**YSD**), we identified Nbs using short synthetic peptides derived from hnRNPA2/B1 and tau. Although the affinities of the identified Nbs were relatively weak, they showed high specificity towards the respective target. We validate their binding properties in cells and demonstrate targeted degradation using Nb-E3 ubiquitin ligase fusions. Directed evolution of the hnRNPA2/B1 binding Nb yielded higher affinity clones, but the mutations caused intracellular aggregation, demonstrating the challenge in engineering Nb affinity while preserving the intracellular stability.

## 2. Methods

### Yeast surface display nanobody libraries

Previously reported YSD Nb libraries were used (a gift from Dr. Andrew Kruse and Dr.

Aashish Manglik) (34, 35). An extensive, detailed step-by-step library screening protocol is described in the **Supplementary Methods**. Briefly, induced YSD Nb libraries were first enriched with antigen-coated magnetic beads by magnetic-activated cell sorting (**MACS**) for 3 rounds. Enriched Nb libraries were then further sorted with fluorescence-activated cell sorting (**FACS**) with multivalent antigens preloaded onto streptavidin. After 4 rounds of FACS sorting, libraries were plated, and individual clones were isolated and sequenced using Sanger sequencing.

### Synthetic peptides and shRNA

Lyophilized hnRNPA2/B1 peptide (SSMAEVDAAMAARPHSIDGRVVEPKK-biotin), tau peptide (RGAAPPGQKGQANATRK-biotin), and three control peptides (C1: KKVAVVRTPPKSPSSAKC-biotin C2: NVKSKIGSTENLKHQPGK-biotin, C3: KTPPAPKTPPSSGEPPK-biotin, and C4: VQIINKKLDLSNVQSKCK-biotin) were purchased from Peptide 2.0 and dissolved in DNase-, RNase-free water (Fisher Scientific, cat. #PB2819-1) before use. For long-term storage, lyophilized peptides were kept at −80 °C.

hnRNPA2/B1 specific shRNA-H1 (<u>CAGAAATACCATACCATCAA</u>TCTCGAGA<u>TTGATGGTATGGTATTTCTG</u>) (36) was cloned under human U6 type III RNA polymerase III promoter by using a long reverse primer overlapping U6 promoter (37). For a complete list of primer sequences used, please refer to **Supp. Table 4**.

### Plasmid generation

E3 ubiquitin ligase and ubiquitin ligase mimics fused to an anti-GFP Nb cAbGFP were cloned under the CMV promoter (38–43). Exact E3 ligase amino acid sequences cloned in this study are provided in **Supp. Table 2**. cAbGFP Nb sequence used in this study is given in **Supp. Table 1** (44). Primers used for cloning are given in **Supp. Table 4**.

mCherry expressing vector was generated by polymerase chain reaction (PCR) amplifying mCherry coding gene with primers flanked by BamHI (NEB) and NotI (NEB) restriction enzymes, and ligated into the pN3 backbone (Addgene #24544) backbone digested with the same set of enzymes.

The EGFP-hnRNPA2/B1 expression plasmid was cloned using Golden Gate Assembly and expressed from the CMV promoter. For generating nuclear export signal (NES) tagged EGFP-hnRNPA2/B1, a long primer encoding a strong nuclear export signal from protein kinase A inhibitor (NSNELALKLAGLDINK) (45) with EGFP overlap was used with hnRNPA2/B1-specific reverse primer. NES-EGFP-hnRNPA2/B1Δ(PY-NLS) and NES-EGFP-hnRNPA2/B1Δ(LCD) were similarly obtained by using appropriate reverse primers specific to hnRNPA2/B1. The list of primers used for this cloning is given in **Supp. Table 4**.

mCherry-tagged Nbs were cloned under the CMV promoter. Briefly, Nb sequences from yeast expression vector were PCR amplified with primers flanked by BamHI (NEB) and SpeI (NEB) restriction enzymes, and mCherry was PCR amplified with primers flanked by SpeI (NEB) and NotI (NEB) restriction enzymes and ligated into the pN3 backbone digested with BamHI and NotI restriction enzymes. The list of primers used for this cloning is given in **Supp. Table 4**. A previously described plasmid for expressing miRFP-tagged Tau (0N4R) was used as a control in some experiments (46).

### Nanobody binding and affinity measurement

The DNA of Nb 19 with stalk protein sequence was digested from pYDS649 v.2 (NTC selection marker) vector using NotI (NEB) and XhoI (NEB) restriction enzymes and cloned into pYDS649 v.1 vector (TRP selection marker) digested with the same restriction enzymes. The tauNb1 gene was amplified by PCR and cloned into the pCT backbone (47). EBY100 cells (a gift from Dr. Eric Shusta) were then transformed with Nb 19 or tauNb1 expressing plasmids. Yeast cells were then induced for 48 hours in SG-CAA media, and antigen binding was then measured at various biotinylated peptide concentrations. Anti-HA mouse primary antibody (BioXcell, cat. #RT0268, 1:1000 dilution) and anti-mouse FITC (Thermo Fisher, cat. #A11001, 1:1000 dilution) or anti-mouse Alexa 647 (Thermo Fisher, cat. #A21236, 1:1000 dilution) secondary antibodies were used for checking Nb display. Streptavidin-PE (Invitrogen, cat. #S866, 1:250 dilution) was used to label biotinylated hnRNPA2/B1 or tau peptide on the yeast surface. Antibody and antigen labeling were carried out in HEPES buffer. Extensive buffer and media compositions are described in the **Supp. Methods**.

### Nanobody affinity maturation

For error-prone PCR, Nb 19 was amplified with (homology2 primers, **Supp. Table 4**) with Taq DNA polymerase, native (Invitrogen cat. # 18-038-018). 50 µL PCR reaction mix composition was: 100 ng of plasmid template, 200 µM of dNTP mixture (Thermo Fisher Scientific), *Taq* PCR buffer (without MgCl_2_), 2 µM 8-oxo-dGTP (TriLink BioTechnologies), 2 µM dPTP (TriLink BioTechnologies), and primers at 0.5 µM concentration. The following PCR cycle was carried out: 90 sec at 94 °C, 24 cycles of [30 sec at 94 °C, 30 sec at 55 °C, 45 sec at 72°C], 10 mins at 72 °C, hold at 12 °C.

For DNA shuffling, yeast libraries were mini-prepped and the Nb genes were PCR amplified with the homology2 primers. ∼ 5 µg of PCR product was digested with the DNase I enzyme (NEB). The following conditions were programmed to digest the PCR product: digestion reaction mix except DNase I was preheated to 12.5 °C for 15 mins, then 0.2 U of DNase I was added to the reaction, digestion was carried out for 2 mins at 12.5 °C, and the digestion reaction was terminated by heating the reaction to 90 °C for 10 mins. ∼200 ng of digestion products were re-assembled without primers with the following PCR conditions: 90 sec at 94 °C, 30 cycles of [30 sec at 94 °C, 45 sec at 55 °C, 90 sec at 72 °C], 10 mins at 72 °C, hold at 12 °C. The band corresponding to approximately ∼450 base pairs was gel-purified and amplified with regular PCR with homology2 primers. PCR amplified fragments were then transformed into *S. cerevisiae* strain EBY100 along with BsrGI/NheI digested pCT4RE plasmid as previously described (47).

The fluorescence-activated cell sorting process for directed evolution is described in the

## Supplementary Methods

### HEK293FT and HEK293-GFP culture and transfection

HEK293FT cells (ThermoFisher cat. # R70007) and HEK293-GFP cell line (Fisher Scientific cat. # 50-672-586) were maintained in DMEM (Fisher Scientific cat. #12320-032) supplemented with 10% heat-inactivated Fetal Bovine Serum (FBS, Fisher Scientific cat. #MT35011CV). Cells used in this study were not passaged more than 15 times. Unless otherwise stated, HEK293 cells were transfected at 70-80% confluence using Lipofectamine^TM^ 3000 transfection reagent (ThermoFisher cat. #L3000001) by following the manufacturer’s recommended protocol.

For flow cytometry analysis, cells were harvested 36 hours post-transfection by trypsinization (Fisher Scientific cat. #MT25051CI). Cells were washed twice with ice-cold PBS buffer before analysis.

### E3 ligase characterization and intracellular protein degradation

HEK293-GFP cells were grown in 24-well plates (tissue culture treated) and, at about ∼80% confluency, were co-transfected with plasmids expressing E3 ligase-cAbGFP fusion and mCherry in order to monitor transfection efficiency. Approximately 400 ng of E3 ligase-cAbGFP expression plasmid and 50 ng of mCherry expression plasmid were used for each well. After 36 hours, cells were collected after trypsinization, washed once, and resuspended in ice-cold PBS buffer. GFP level in mCherry-expressing cells was then analyzed by using flow cytometry. Cell line maintenance and transfection reagents used are described above.

For intracellular targeted degradation of hnRNPA2/B1, HEK293FT cells were co-transfected with plasmids expressing EGFP-hnRNPA2, E3 ligase-Nb fusion, and mCherry, in 3:16:1 ratio (75 ng:400 ng:25 ng). Cells were harvested 36 hours post-transfection as described above and GFP levels were measured in mCherry-expressing cells using flow cytometry.

### Colocalization analysis

Nb19 was fused C-terminally to mCherry (Nb19-mCherry) and expressed in HEK293FT or HeLa cells using the CMV promoter. EGFP fused to hnRNPA2 (EGFP-hnRNPA2) or hnRNPA2 with an added nuclear export signal (NES) and a deletion of the nuclear targeting sequence (NTS) (EGFP-NES-hnRNPA2ΔNTS) were also expressed using the CMV promoter. A rabbit polyclonal anti-hnRNPA2/B1 antibody (ThermoFisher cat. # PA5-34939, RRID AB_2552288) was used to stain endogenous hnRNPA2/B1.

Confocal microscopy images were taken using either the Nikon A1R or Nikon AXR confocal microscope (University of Connecticut COR^2^E facility). Pearson’s correlation coefficient (PCC, R value above threshold) was calculated using the Coloc 2 plugin in ImageJ by manually selecting regions positive for Nb fluorescence. For each condition, PCC was calculated from 5-6 regions taken from 3-4 images. The mean and standard deviation of PCCs are reported.

### Flow Cytometry

Flow cytometry data were obtained using a BD Biosciences LSRFortessa X-20 Cell Analyzer (University of Connecticut COR^2^E facility). Fluorescence-activated cell sorting was carried out using the BD Aria II Cell Sorter (University of Connecticut COR^2^E facility). Data were analyzed using FlowJo (vX.0.7).

## 3. Results

### Yeast display screening of an anti-hnRNPA2/B1 nanobody

hnRNPA2/B1 has four isoforms named A2, B1, A2b and B1b due to alternative splicing (4). All isoforms have N-terminal tandem RNA-binding domains, RNA-recognition motif 1 (**RRM1**) and RRM2, and a C-terminal glycine-rich low complexity region (4) (**Fig. 1a**). Isoforms A2 and B1 are often referred to together as hnRNPA2/B1, since they differ only by a 12-amino acid insertion at the N-terminus (see **Supp. Fig. 1** for an alignment of A2 and B1 isoforms). To screen Nb libraries, we used a peptide sequence derived from the RRM1 domain (**Fig. 1a, Supp. Fig. 1**). To increase the efficiency of identifying potentially weak Nb binders, we employed a screening approach that leverages the avidity of binders displayed on the yeast surface. In the first screening stage, we incubated magnetic beads pre-loaded with a peptide sequence derived from hnRNPA2/B1 with yeast display Nb libraries to enrich for binders using magnetic-activated cell sorting (**MACS**) (**Fig. 1b**). In the second stage, we conducted fluorescence-activated cell sorting (**FACS**) using peptides pre-loaded with fluorescently labeled streptavidin (**Fig. 1c**). For the antigen-coated magnetic bead preparation, the optimal antigen to magnetic bead ratio was determined empirically as described in **Supp. Methods** (**Supp. Fig. 2**). We kept the peptide-to-magnetic bead ratio constant, but lowered the concentration of the preloaded magnetic beads as the MACS stage progressed (**Fig. 1d**). We also performed depletion steps with magnetic beads without antigen to remove non-antigen binders (**Fig. 1d**). After MACS, we conducted four rounds of FACS to obtain a final enriched pool (**Fig. 1d**). We observed substantial enrichment of antigen binders while the fraction of streptavidin-PE binders remained minimal (**Fig. 1e**).

**Figure 1.**
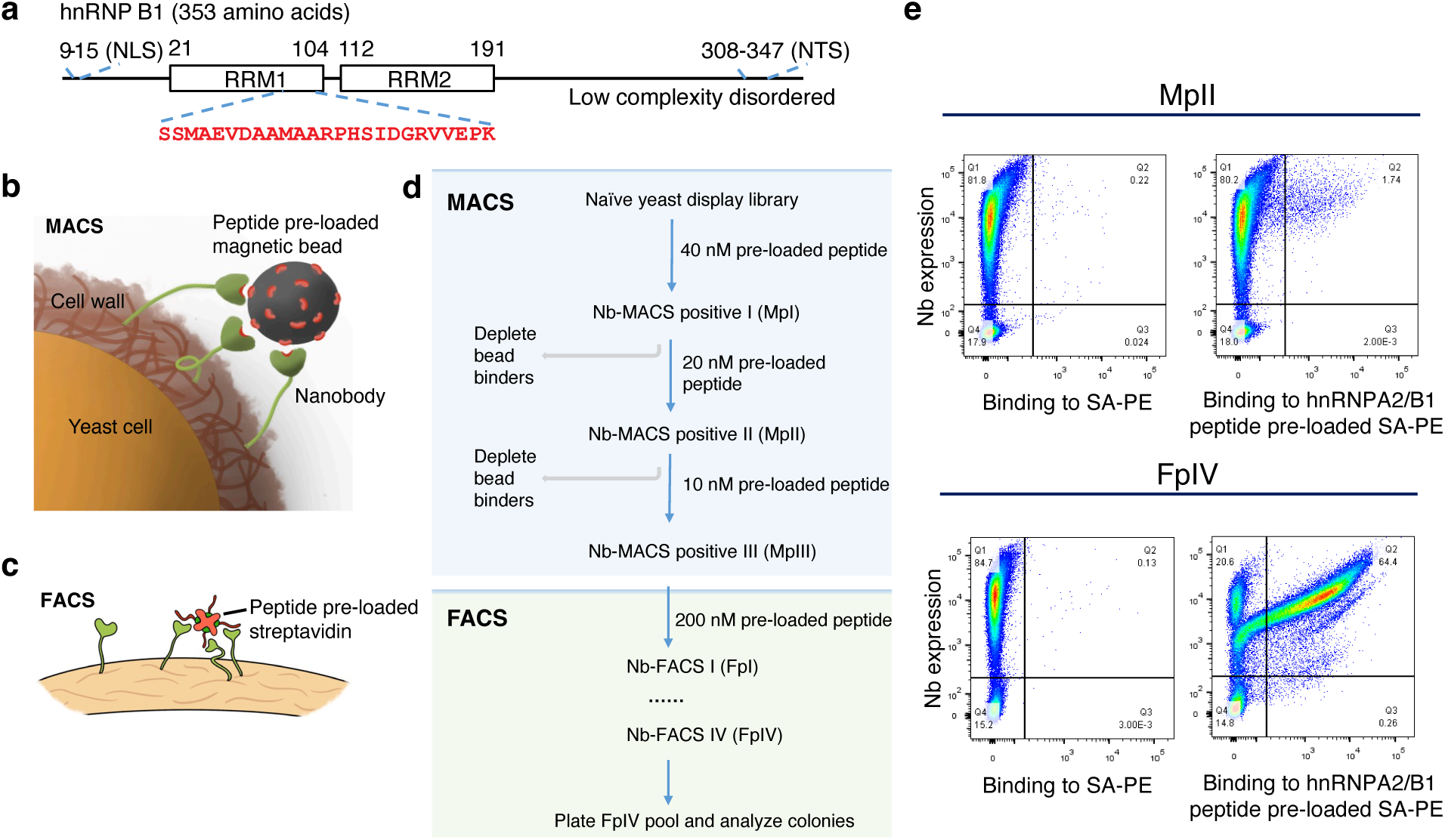
Nb library screening against hnRNPA2/B1 enabled by a multivalent library screening approach. **(a)** Schematic of the hnRNP A2/B1 sequence. The target peptide sequence used for the screen is indicated in red. NLS – nuclear localization signal, NTS – nuclear targeting sequence. **(b)** Streptavidin-coated magnetic beads pre-loaded with the target peptide were used for MACS. **(c)** The target peptide pre-loaded with streptavidin was used for FACS. **(d)** Screening flow chart for enriching hnRNPA2/B1 Nbs. (**e**) Enrichment of binder clones during the MACS and FACS stages. The pool of clones after indicated rounds of screening was labeled with the target peptide preloaded Streptavidin-PE (SAPE) or SAPE only. Pre-loaded peptide concentration was 400 nM.

**Figure 2.**
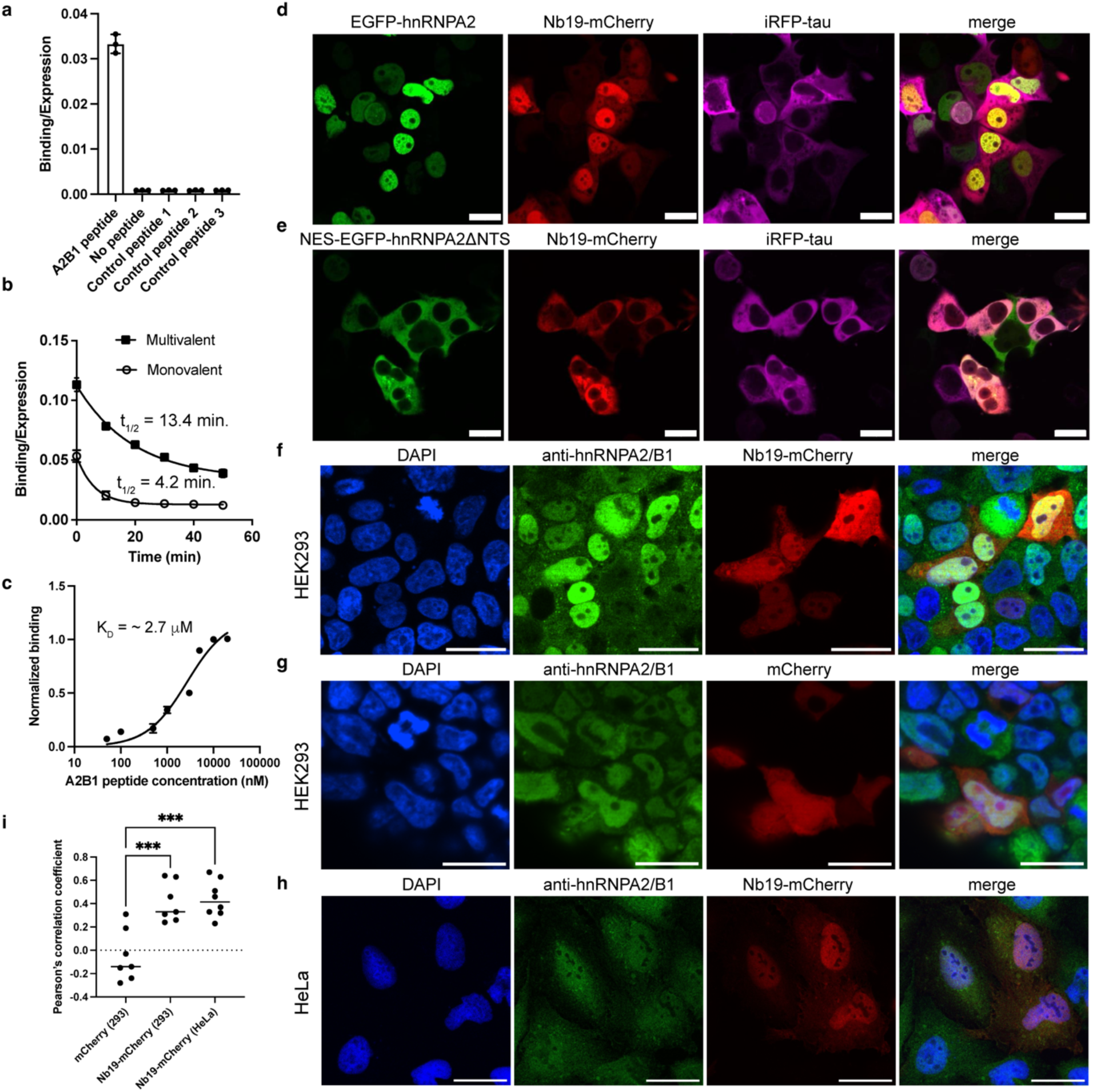
Nb characterization. **(a)** Yeast cells displaying Nb19 were incubated with 1 µM biotinylated hnRNPA2/B1 peptide (A2B1) or control peptides. No peptide indicates cells incubated with SAPE without any peptide. **(b)** Yeast cells displaying Nb19 were labeled with 10 nM A2B1 with or without SAPE pre-loading. After incubation, excess peptide was removed, and cells were resuspended in buffer and left at room temperature. Fluorescence from yeast cells was measured every 10 minutes for 50 minutes. **(c)** Yeast cells displaying Nb 19 were incubated with varying concentrations of A2B1. In panels **(a-c)** fluorescence was measured using flow cytometry. **(d, e)** Confocal microscopy image of HEK293FT cells expressing Nb19-mCherry, miRFP fused tau, and **(d)** EGFP-hnRNPA2 or **(e)** NES-EGFP-hnRNPA2ΔNTS. In NES-EGFP-hnRNPA2ΔNTS, a nuclear-export signal (NES) from protein kinase A inhibitor was fused to the N-terminus, and the NTS was deleted. In addition, in panel **(e)**, shRNA specific to only endogenous hnRNPA2/B1, but not to NES-EGFP-hnRNPA2ΔNTS, was co-expressed. (**f, g**) Confocal microscopy images of HEK293FT cells stained with an anti-hnRNPA2/B1 antibody expressing **(f)** Nb19-mCherry or **(g)** mCherry. **(h)** The same as in panel (f) but in HeLa cells. **(i)** Pearson’s correlation coefficient (R value) calculated from confocal microscope images (7-8 regions across 4-5 images) of HEK293FT cells and HeLa cells. ***, *P* < 0.001 Dunnett’s multiple comparisons test. Plotted in panels (**a-c**) are mean and standard deviation from 3 biological replicates. In panels (**d-h**), scale bars indicate 20 μm.

### hnRNPA2/B1 Nb characterization

From the enriched pool of binders, we selected a Nb clone we named Nb 19 for further analysis (see **Supp. Table 1** for Nb sequence). Yeast cells displaying Nb 19 showed clear binding to streptavidin pre-loaded with the hnRNPA2/B1 peptide used for screening, but not to bare streptavidin or streptavidin pre-loaded with other peptides (**Fig. 2a**). The control peptide sequences are derived from the human tau protein. To assess the effect of peptide pre-incubation, we compared the dissociation rate of free peptide and peptides pre-loaded with streptavidin (**Fig. 2b**). The half-lives of dissociation of free peptides (monovalent) and pre-loaded peptides (multivalent) were 4.2 min. and 13.4 min., respectively (**Fig. 2b**). This shows that the avidity gained by streptavidin pre-loading slows the dissociation rate. We measured the monovalent binding affinity of Nb19 by displaying it on yeast and incubating the yeast cells with various concentrations of the hnRNPA2/B1 peptide. The dissociation constant (K_d_) was 2.7 μM with a 95% confidence interval of 1.9 to 3.8 μM (**Fig. 2c**). These results show that Nb19 binds to the target epitope in hnRNPA2/B1 with a micromolar affinity when expressed on the yeast surface.

We also tested Nb19 binding to hnRNPA2/B1 when expressed intracellularly. We expressed Nb19 fused to mCherry at the C-terminus (Nb19-mCherry) in HEK293FT cells. We co-expressed hnRNPA2 N-terminally fused to EGFP (EGFP-hnRNPA2) and assessed colocalization with Nb19-mCherry. Nb19-mCherry fluorescence showed a positive correlation with EGFP-hnRNPA2 (**Fig. 2d**, Pearson’s R = 0.51 ± 0.16). To further test the intracellular binding, we expressed EGFP-hnRNPA2 in the cytosol by fusing a nuclear export signal (NES) and deleting the nuclear targeting sequence (NTS) (EGFP-NES-hnRNPA2ΔNTS). In this experiment, we also co-expressed a shRNA specific to endogenous hnRNPA2/B1, but not to ectopically expressed EGFP-hnRNPA2/B1. Nb19-mCherry fluorescence shifted to the cytosol with a positive correlation to EGFP-NES-hnRNPA2ΔNTS (**Fig. 2e**, Pearson’s R = 0.70 ± 0.13). Although EGFP-hnRNPA2 is mostly localized to the nucleus, Nb-mCherry fluorescence is also detected in the cytoplasm (**Fig. 2d**). This may partly be due to the overexpression of the Nb, but also can be attributed to spliceoforms A2b and B1b localized in the cytoplasm (48) (see **Discussion**). To test this, we assessed colocalization of Nb19-mCherry with an anti-hnRNPA2/B1 antibody staining. In HEK293FT cells, Nb19-mCherry showed significantly higher positive colocalization with the antibody compared to mCherry without the Nb (**Fig. 2f, g, i**). Nb19-mCherry also showed significant positive colocalization in HeLa cells (**Fig. 2h, i**). The antibody staining was stronger in the nucleus but also detected in the cytoplasm (**Fig. 2g, i**), indicating that a fraction of endogenous hnRNPA2/B1 is in the cytoplasm. Taken together, we conclude that Nb19 binds to the target epitope in hnRNPA2/B1 and binds to full-length hnRNPA2/B1 when expressed intracellularly.

### Yeast display screening of an anti-tau Nb

Given the interaction of hnRNPA2/B1 and the microtubule-associated protein tau, we also set out to screen the Nb library against tau. We selected a region of the tau protein that shows weak homology between human and mouse tau (**Fig. 3a**, **Supp. Fig. 4**), which is the epitope targeted by the human tau-specific antibody HT7. Using the same Nb screening pipeline, but with the tau peptide, we identified a Nb that we named tauNb1 (**Supp. Table 1**). Yeast cells displaying tauNb1 showed binding to streptavidin pre-loaded with the target tau peptide, but not to streptavidin pre-loaded with other peptides also derived from the human tau protein (**Fig. 3b**). Although we clearly detected the binding of streptavidin pre-loaded with the target tau peptide (**Fig. 3c**, multivalent), we did not detect significant monovalent binding of peptides (**Fig. 3c**, monovalent), even at 10 μM peptide concentration. This suggests that the K_d_ of tauNb1 is much larger than 10 μM. Even though the affinity is low, when we expressed tauNb1-mCherry with EGFP-tau in HEK293FT cells, we observed a strong positive colocalization (**Fig. 3d**, Pearson’s R = 0.74 ± 0.10, n = 6 regions from 5 images). When we expressed EGFP-tau fused to a strong nuclear localization signal (NLS-EGFP-tau) and co-expressed tauNb1-mCherry, we also observed a strong positive colocalization of EGFP and mCherry fluorescence (**Fig. 3e**, Pearson’s R = 0.73 ± 0.15, n = 6 regions from 5 images). These results show that tauNb1 has low affinity, but can bind to the tau protein when expressed in cells.

**Figure 3.**
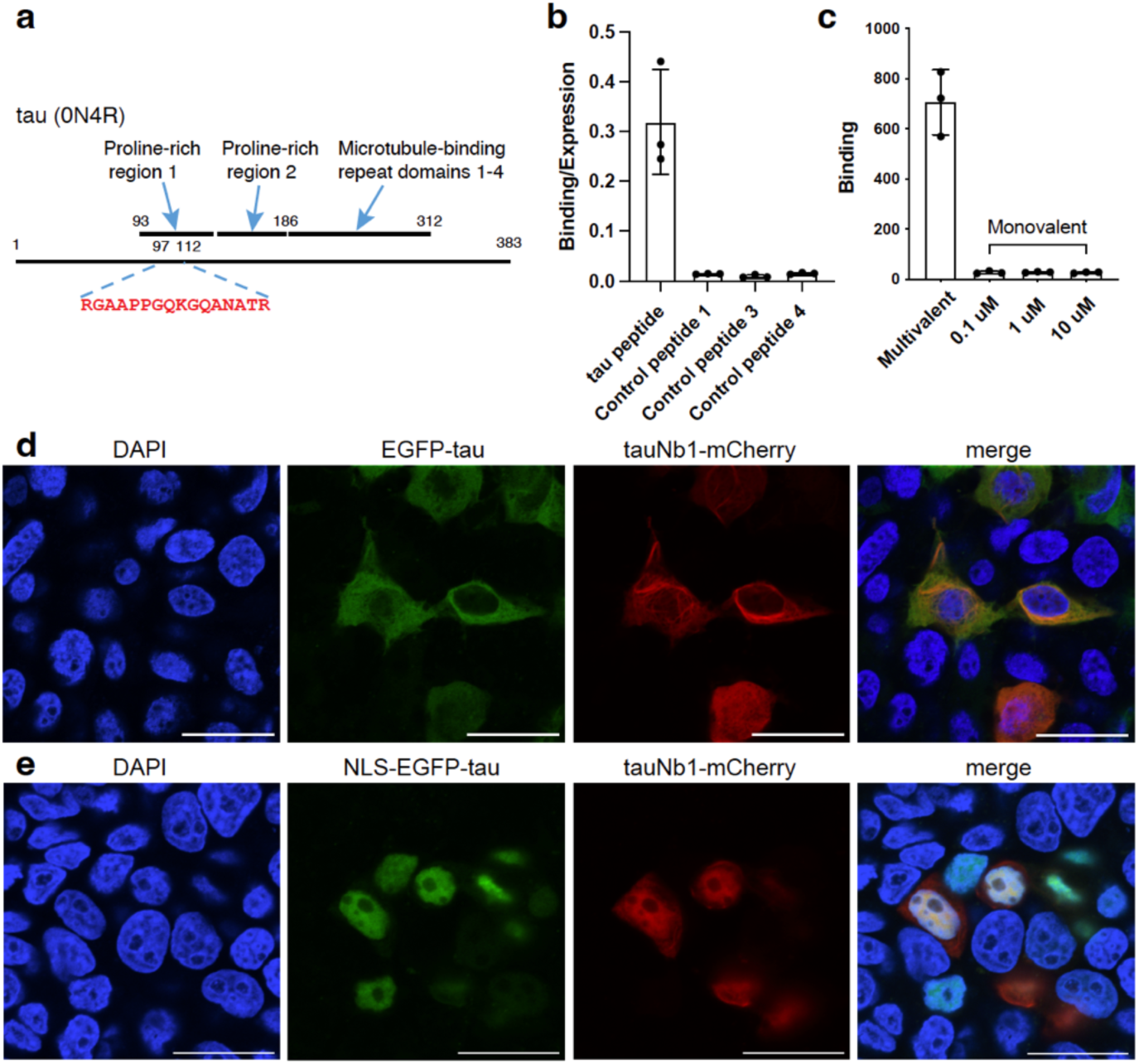
Anti-tau Nb tauNb1. **(a)** Schematic of the human tau (0N4R isoform) sequence. The target peptide sequence used for the screen is indicated in red. PRR – proline-rich region, MTBR – microtubule binding region. **(b, c)** Confocal microscopy image of HEK293FT cells expressing tauNb1-mCherry and **(b)** EGFP-tau (0N4R) and **(c)** EGFP-tau (0N4R) fused to the NLS from Myc. Scale bars indicate 25 μm.

**Figure 4.**
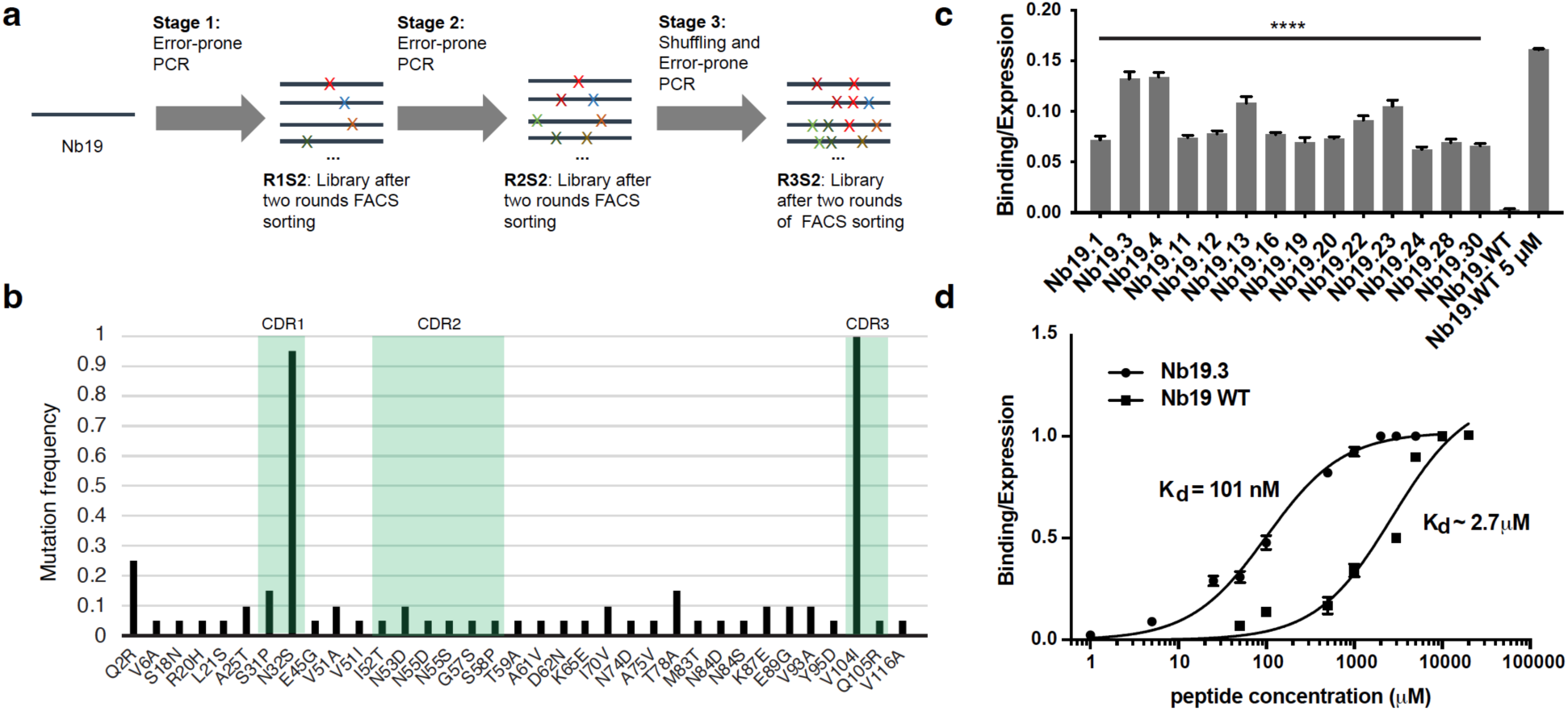
Directed evolution of Nb19. **(a)** Overview of directed evolution stages. In stage 1, error-prone PCR was used to introduce mutations, and improved binding clones were selected using two rounds of FACS, resulting in pool R1S2. Stage 1 was repeated in stage 2, leading to pool R2S2. In stage 3, DNA shuffling was used to combine the mutations, followed by another error-prone PCR step. Finally, clones with higher binding were selected using two rounds of FACS, leading to pool R3S2. **(b)** Mutation frequency analysis in selected clones. A total of 20 unique clones were sequenced, and mutation frequencies were calculated. Mutations in CDRs are highlighted in green. **(c)** Yeast cells displaying Nb clones were labeled with 15 nM of hnRNPA2/B1 peptide and binding/expression (display) was calculated. Wild-type (WT) Nb19 was incubated with 15 nM or 5 µM hnRNPA2/B1 peptide. Plotted are the means and standard deviation from 3 biological replicates. ****, *P* < 0.0001 Dunnett’s multiple comparisons test. **(d)** Binding/expression of Nb19.3 to varying hnRNPA2/B1 peptide concentrations. Plotted are means and standard deviations from 3 biological replicates. Data for WT Nb19 were replotted from Figure 1c for comparison.

### Directed evolution of anti-hnRNPA2/B1 Nb

We next pursued improving the Nb affinity through directed evolution. To increase the affinity of Nb19, we carried out three stages of directed evolution using YSD (**Fig. 4a**). In the first stage, we performed error-prone PCR of the Nb19 gene and screened for clones with improved binding using two rounds of fluorescence-activated cell sorting (**FACS**). In the second stage, we induced further mutations using error-prone PCR, then sorted for improved binders using two rounds of FACS. Finally, in the third stage, we shuffled the isolated Nb genes, then conducted error-prone PCR. After two rounds of FACS, we randomly selected 30 clones from the isolated pool and checked their antigen binding with 20 nM (monovalent) of hnRNPA2/B1 peptide. 23 clones from the 30 clones showed substantial binding, which were sequenced. From this, a total of 20 unique Nb sequences were obtained (**Supp. Table 3**), and the frequency of each mutation was calculated (**Fig. 4b**). This showed that the mutations N32S and V104I are observed in >90% of clones, suggesting that the selection pressure strongly favored these mutations. The mutations N32S and V104I are located in complementarity-determining regions (**CDRs**) 1 and 3, respectively (**Fig. 4b**). From the 20 clones, we selected 14 and incubated them with 15 nM (monovalent) hnRNPA2/B1 peptide. For the 14 mutant clones, the binding signal normalized to the Nb expression level was significant at 15 nM peptide, while the original Nb19 had no detectable signal above background (**Fig. 4c**). We further characterized the affinity of Nb19.3, which showed strong binding at 15 nM. Using titration of the hnRNPA2/B1 peptide, we observed K_d_ = 101 nM for Nb19.3 (**Fig. 4d**), which represents a 27-fold increase in binding affinity. These results show that improved affinity mutants were successfully selected.

### Higher-affinity Nb mutant aggregates in the cytoplasm

We next characterized the higher-affinity mutant Nb19.3 in HEK293FT cells. When we expressed Nb19.3 fused to mCherry (Nb19.3-mCherry), we observed puncta in the cytoplasm that were not seen with Nb19 (**Fig. 5a**). For HEK293FT cells expressing Nb19-mCherry or Nb19.3-mCherry, we quantified the percentage of cells with visible Nb puncta. Among Nb expressing cells (mCherry positive), we observed puncta in 2.7 ± 1.4 % of cells for Nb19, compared to 92.5 ± 3.9 % of cells for Nb19.3 (**Fig. 5b**). When we stained Nb19.3-mCherry expressing cells with an anti-hnRNPA2/B1 antibody, the Nb puncta did not colocalize with antibody staining (**Fig. 5c, d**). To check if the Nb19.3 puncta colocalize with lipid-bound vesicles, we stained the cells with a lipid droplet dye (Biotracker 488). The dye stain did not colocalize with the puncta (**Fig. 5e, f**). To test whether Nb19.3 puncta are dynamic and capable of exchanging proteins, we measured fluorescence recovery after photobleaching (**FRAP**) of Nb19.3-mCherry. Fluorescence from Nb19.3-mCherry puncta did not recover after photobleaching (**Fig. 5g, i**), while fluorescence from diffuse Nb19.3-mCherry recovered to 82% of the initial value (**Fig. 5h, i**). This result shows that the Nb19.3 forms stable aggregates in the cytoplasm. When we mutated the residues in Nb19.3 back to the Nb19 sequence, we found that the fraction of cells with the puncta decreased significantly for mutations 32N and 104V (**Fig. 5b**), which are located in CDR1 and CDR3, respectively. The 100V mutation within framework 3 did not change the fraction of cells with the puncta significantly (**Fig. 5b**). Overall, the puncta formation was not attributed to a single mutation. We also introduced a mutation (45G) that was previously reported to improve Nb stability (49). The mutation reduced puncta formation significantly, but only by about 20% (**Fig. 5b**). These results show that the mutations that increased binding affinity caused the formation of stable aggregates in the cytoplasm.

**Figure 5.**
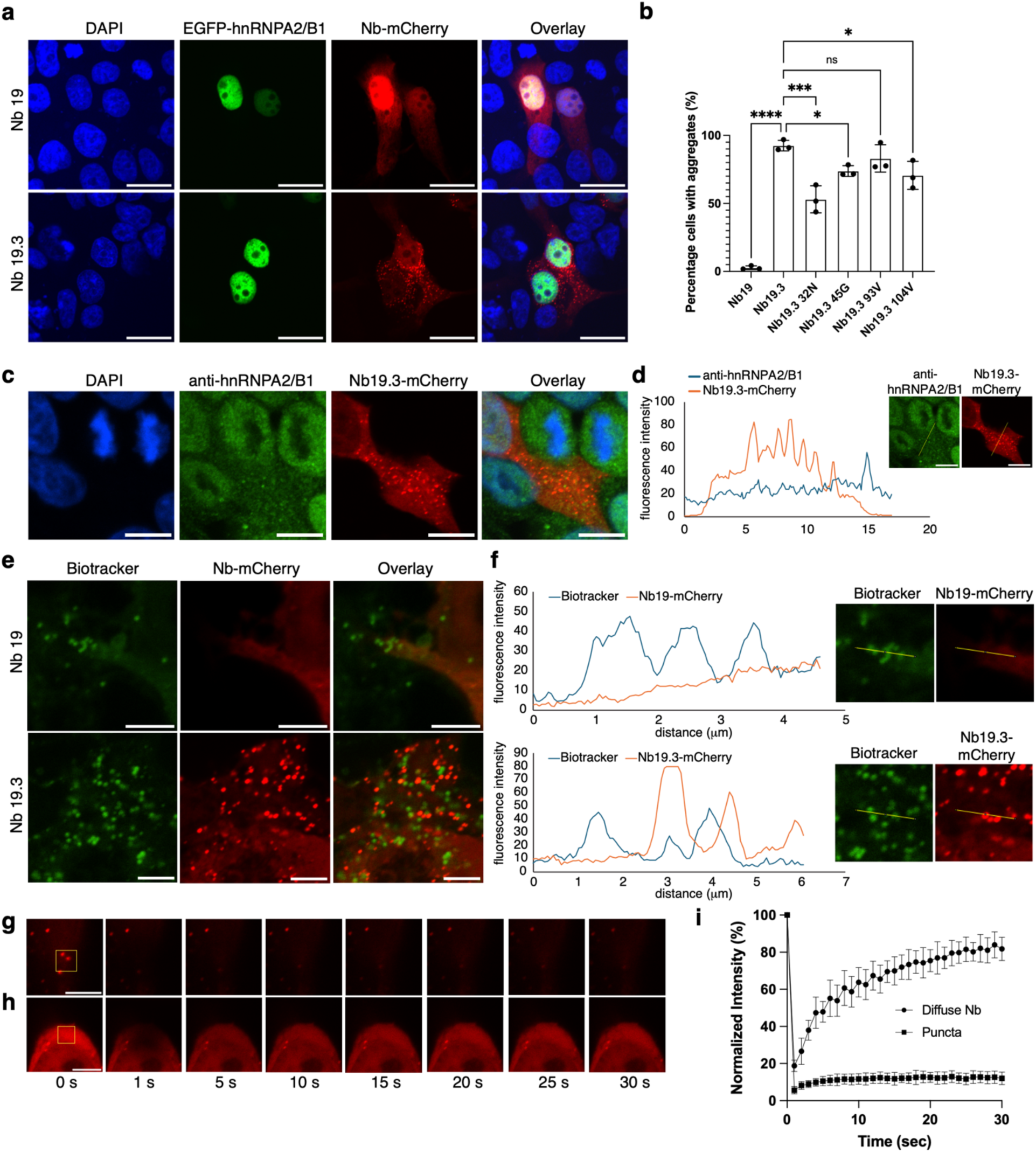
Aggregation of Nb19.3 in the cytoplasm. **(a)** Confocal microscopy images of HEK293FT cells co-expressing EGFP-hnRNPA2, and Nb19-mCherry or Nb19.3-mCherry. Scale bars indicate 20 μm. **(b)** Percentage of aggregation-positive HEK293FT cells expressing mCherry fusions of the indicated Nbs. Plotted are means and standard deviation from 3 biological replicates. *, *P* < 0.05; ***, *P* < 0.001; ****, *P* < 0.0001 Dunnett’s multiple comparisons test. **(c)** Confocal microscopy images of HEK293FT cells expressing Nb19.3-mCherry stained with an anti-hnRNPA2/B1 antibody. Scale bars indicate 10 μm. **(d)** Fluorescence intensity profile of anti-hnRNPA2/B1 antibody stain and Nb19.3-mCherry. Insets show the lines used to generate the profiles. **(e)** Confocal microscopy images of HEK293FT cells expressing Nb19.3-mCherry stained with the Biotracker 488 lipid droplet dye. Scale bars indicate 5 μm. **(f)** Fluorescence intensity profile of the Biotracker stain and Nb19.3-mCherry. Insets show the lines used to generate the profiles. (**g, h**) Fluorescence recovery after photobleaching (FRAP) of Nb19.3-mCherry puncta (**g**) and diffuse Nb19.3-mCherry (**h**). Scale bars indicate 5 μm. Yellow boxes indicate photobleached regions. (**i**) Fluorescence measurement during FRAP of Nb19.3-mCherry puncta and diffuse Nb19.3-mCherry. Plotted are the means and standard deviation from 3 biological replicates.

### Targeted degradation using Nb-E3 ligase fusions

To further demonstrate the use, we tested Nb-mediated targeted degradation of hnRNPA2/B1 and tau proteins. A general design principle is to fuse Nbs with an E3-ligase adapter domain that recruits the ubiquitin proteasome system for degradation of the Nb-bound protein. We first characterized previously known and several new E3 ligase adapters and E3 ligase mimics for targeted protein degradation in HEK293 cells. We constructed a panel of Nb-E3 ligase fusions by replacing the substrate recognition domains of these proteins with a GFP-binding Nb (see **Supp. Table 2** for Nb-E3 fusion sequences). The synthetic E3 ligase constructs were transfected into a GFP-expressing HEK293 stable cell line, along with mCherry as a transfection control. GFP levels in mCherry-positive cells were measured using flow cytometry (**Fig. 6a, Supp. Fig. 3**). We observed a substantial decrease in GFP level in cells expressing the previously reported synthetic E3 ligase mimic IpaH9.8 (40) (**Fig. 6a**). Based on this information, we used IpaH9.8 as the E3 ligase adaptor in all subsequent experiments. To detect changes to the hnRNPA2 or tau level, we generated a plasmid that expresses miRFP and EGFP-hnRNPA2 (or EGFP-tau) using the ribosome skipping sequence P2A (**Fig. 6b, c**). We then co-expressed the P2A construct and Nb-IpaH9.8 fusions (**Fig. 6b, c**) in HEK293FT cells and measured the miRFP and EGFP fluorescence using flow cytometry. The EGFP fluorescence normalized to the miRFP fluorescence was reduced by 69% in cAbGFP4-IpaH9.8 co-expressing cells and by 41% in Nb19-IpaH9.8 co-expressing cells compared to those co-expressing the control Nb-IpaH9.8 (**Fig. 6d**). Nb19.3-IpaH9.8 also showed similar reduction in EGFP-hnRNPA2 levels (**Fig. 6d**). We also tested the tauNb1 for targeted degradation of EGFP-tau (**Fig. 6c**) and found that the EGFP/miRFP ratio was reduced by 46% in cells co-expressing cAbGFP4-IpaH9.8 and 34% in cells co-expressing tauNb1-IpaH9.8 compared to those co-expressing the control Nb-IpaH9.8 (**Fig. 6e**). These results indicate that Nb19, Nb19.3, and tauNb1 can direct IpaH9.8 for targeted degradation, although Nb cAbGFP4 was more efficient. We also tested if endogenous hnRNPA2/B1 can be degraded using Nb-IpaH9.8 fusions. We transfected plasmids encoding Nb19-mCherry, cAbGFP4-IpaH9.8, Nb19-IpaH9.8, or Nb19.3-IpaH9.8 into HEK293T cells, and probed hnRNPA2/B1 using Western blotting. Compared to cells expressing Nb19-mCherry, hnRNPA2/B1 levels (normalized to GAPDH) were reduced by 75% in cells expressing Nb19-IpaH9.8 (**Fig. 6f, g**). No significant change was observed in cells expressing cAbGFP4-IpaH9.8 and Nb19.3-IpaH9.8 (**Fig. 6f, g**). These results show that while Nb19-IpaH9.8 significantly lowers the level of endogenous hnRNPA2/B1, Nb19.3-IpaH9.8 is not effective. Taken together, while these results validate the binding activities of Nbs developed in this study, they also illustrate the challenge in engineering improved-affinity Nbs.

**Figure 6.**
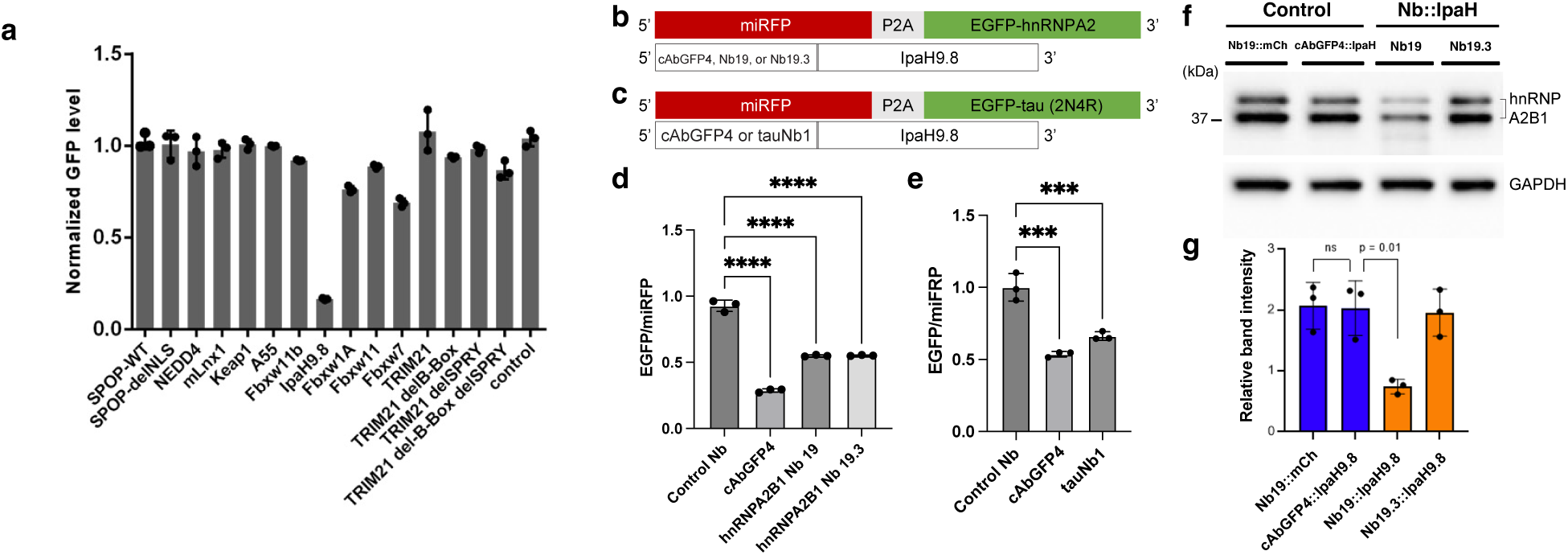
Targeted degradation using Nb-E3 ligase fusions. **(a)** Characterization of ubiquitin E3 ligase domains and E3 mimics for targeted degradation. A panel of E3 ligase domains was fused to the C-terminus of the anti-GFP Nb cAbGFP4. cAbGFP4-E3 ligase domain fusion constructs were transfected into an EGFP-expressing HEK293 cell line along with an mCherry-expressing plasmid. The EGFP levels in mCherry-positive cells were normalized to those of non-transfected cells. Plotted are the mean and standard deviation from three independent experiments. **(b)** Schematic of gene constructs expressing miRFP-P2A-EGFP-hnRNPA2 and Nb-IpaH9.8 fusions. **(c)** Schematic of gene constructs expressing miRFP-P2A-EGFP-tau and Nb-IpaH9.8 fusions. **(d, e)** EGFP level normalized to miRFP in cells co-transfected with the constructs shown in panels (b) and (c), respectively. Plotted are mean and standard deviation from three independent experiments. ****, *P* < 0.0001; ***, *P* < 0.001 Dunnett’s multiple comparisons test. **(f)** Western blots of HEK293T cell lysates transfected with the indicated constructs. **(g)** Quantification of results in panel (f). hnRNPA2/B1 band intensity was normalized to that of GAPDH. Plotted are the mean and standard deviation from three blots. P values are calculated using Student’s t-test.

## 4. Discussion

Here, we report the identification and validation of Nbs binding to human hnRNPA2/B1 and tau. To find the Nbs, we developed an approach to screen YSD libraries using synthetic peptides as antigens. This approach relies on multivalent peptide-Nb interactions for efficient library screening in both MACS and FACS stages (**Fig. 1b, c**). During MACS, multivalent binding was achieved by pre-loading the peptides with streptavidin-coated magnetic beads (**Fig. 1b**). Previous studies have used biotinylated small molecules, peptides, or proteins for screening YSD libraries (34, 50–52). We systematically tested the peptide preloading conditions that allowed for the efficient recovery of peptide-binding yeast cells (**Supp. Fig. 2b**). This multivalent antigen-Nb interaction enhances the apparent binding affinity and enables the efficient labeling of YSD Nb libraries with magnetic beads. Using the multivalent nature of streptavidin itself (without relying on bead immobilization) to generate multivalent antigens (**Fig. 1c**) for FACS screening of YSD libraries has been suggested previously (51); however, to our knowledge, a concrete example of its use in binder screening has not been reported. Our data show that streptavidin preloading reduces the apparent dissociation rate of peptides from the yeast surface (**Fig. 2b**) and enables detecting the binding of a weak-affinity Nb, not detected using monovalent peptide incubation (**Fig. 3c**). This multivalent screening approach may apply to identifying other binders with weak affinity.

A major advantage of using peptides for Nb screening is that a desired epitope can be targeted. In many Nbs, the epitope plays a critical role in their biological function and mechanism of action. For example, in targeting GPCRs, Nbs with different epitopes can stabilize distinct conformations of the same GPCR for structural characterization (53) and modulate signaling differently (29). When neutralizing viruses, targeting distinct epitopes provides alternative mechanisms for neutralization (54, 55). In intracellular targeting of proteins, selecting epitopes that do not interfere with native complex formation can be critical (56). In addition, having non-overlapping epitopes on a single target is required in immunoassay development (57). However, obtaining Nbs against a pre-determined binding site remains a challenge. Since short peptides are weak immunogens, camelid immunization with short peptides generally relies on carrier conjugates (58, 59). Since this may lead to carrier immunodominance (60–62), a large number of clones need to be screened using a secondary assay to find target binders. As for *in vitro* Nb library screening, competition-based selections are used to identify clones that bind to a desired epitope (63–65). Given these drawbacks, being able to screen Nb libraries against a defined epitope is a significant advantage.

Using peptides as antigens has great potential to enable identifying anti-tau Nbs specific to post-translational modifications (PTMs). Peptides containing defined PTMs can be readily synthesized and are widely used for producing PTM-specific anti-tau antibodies (66). Several groups have developed anti-tau Nbs (67–71), but PTM-specific Nbs have not been reported yet. Given that tau PTM is a key mechanism in studying tau pathology (72–74), PTM-specific anti-tau Nbs will be highly valuable. E3 ligase fusions to such Nbs would enable targeted degradation of post-translationally modified tau.

Using Nbs for imaging hnRNPA2/B1 may provide unique advantages in studying its dynamics. When we expressed EGFP-hnRNPA2, we observed that it is almost entirely localized in the nucleus (**Fig. 2d**). The pre-mRNA of hnRNPA2/B1 is spliced to form four isoforms: B1 (includes all exons), A2 (missing exon 2), B1b (missing exon 9), and A2b (missing exons 2 and 9) (75). Although the most abundant A2 isoform is primarily located in the nucleus, a significant fraction of A2b and B1b isoforms is in the cytoplasm, depending on the cell type and developmental stages (48). We also detect hnRNPA2/B1 antibody staining in the cytoplasm (**Fig. 2f, h, and 5c**). Moreover, hnRNPA2/B1 is actively transported in and out of the nucleus (76), and the misregulation of the nucleocytoplasmic transport is associated with various diseases (17, 18, 77, 78). This result illustrates the complexity of imaging endogenous proteins and the advantage that Nbs may have over fluorescent protein fusions.

Although we identified mutations in Nb19 that improve its binding affinity, the mutations caused aggregations when the Nb was expressed in the cytoplasm. This was particularly important for this study, since a major goal of using the nanobodies is to study protein condensates. We therefore extensively characterized the puncta formed in hnRNPA2/B1 nanobodies. A recent study that tested 75 Nbs from the Protein Data Bank in the cytoplasm of mammalian cells found that 42 out of 75 formed intracellular puncta or lacked expression (49). Based on the sequence of stable Nbs, several destabilizing framework residues were identified (49), which were not found in Nb19 or Nb19.3. Also, the same study identified framework residues enriched in stable Nbs (49), most of which were present in Nb19 and Nb19.3, except for the 45G residue (**Fig. 5b**). Introduction of the 45G mutation into Nb19.3 markedly reduced the fraction of cells displaying puncta, although most cells still exhibited them (**Fig. 5b**). Nb19.3 has mutations N32S and V104I, which are present in nearly all affinity improved clones found through YSD screening (**Fig. 4b**). N32S is in CDR1 while V104I is in CDR3. Both of these mutations have a substantial impact on intracellular puncta formation (**Fig. 5b**). Previous studies using thermal denaturation have shown that mutations in the CDR of Nbs can affect stability (79), particularly when the CDR residue interacts with the framework region (80). The reduced cytoplasmic stability of Nb19.3 is likely because the mutant screening was conducted under non-reducing buffer conditions, while the intracellular environment is strongly reducing (81), cleaving the disulfide bond within the Nb framework region (82). Moreover, molecular crowding caused by the excluded volume effect in the cytoplasm can promote aggregation and non-specific binding (83–85). These results demonstrate the need for screening approaches that can improve Nb affinity in the intracellular environment.

We anticipate that the Nbs against the RBP hnRNPA2/B1 and tau, as well as the E3 ligase fusions, will be a useful tool for imaging their dynamics and phase separation behavior. Although the high-affinity mutant Nb19.3 aggregated in the cytoplasm, the Nb may be useful as a binding reagent under non-reducing conditions.

## Acknowledgements

This work was supported by National Institutes of Health (NIH) grants R21 AG083761 and R01 AG083876 (to Y.C.) and R01 AG080810 (to B.W.). The content is solely the responsibility of the authors and does not necessarily represent the official views of the National Institutes of Health.

## Declaration of interests

B.W. is CSO and Co-Founder of Aquinnah Pharmaceuticals Inc.. The other authors declare that they have no conflicts of interest with the contents of this article.

## Author Contributions

A.P., B.W., and Y.C conceived the project; B.W. and Y.C. acquired funding; A.P., B.W., and Y.C. designed research; A.P. conducted the screening experiments with support from C.R., R.S., and M.A.; A.P., Y.C., F.T.A., R.G., K.T., E.E., S.P. characterized the nanobodies; A.P., C.R., F.T.A., R.G., B.W., and Y.C. analyzed data; A.P. and Y.C. wrote the manuscript; all authors edited the manuscript.

