## Supplementary Materials for "Nanobodies targeting hnRNPA2/B1 and tau"

### Supplementary Methods

#### a. Determining optimal magnetic bead to peptide ratio for magnetic bead preloading

The optimal magnetic bead to peptide ratio for generating peptide-preloaded magnetic beads was empirically determined. We used previously published yeast cells [1] displaying a control scFv 4420 (also expressing mCherry in the cytoplasm) or an anti-phospho-tau scFv 3.24 (binds to peptide-peptide KKVAVVRpTPPKpSPSSAKC, express EGFP in the cytoplasm). Briefly, approximately $6 \times 10^{16}$ to $6 \times 10^{6}$ biotinylated-peptides (KKVAVVRpTPPKpSPSSAKC-biotin) were incubated with 10 µL of streptavidin microbeads (Miltenyi Biotec, Cat. No. 130-048-012) in 30 µL PBS buffer for 1 hour at room temperature with constant mixing. Induced yeast cells displaying scFv 4420 or scFv 3.24 were mixed in equal numbers. A total of $2 \times 10^{8}$ cells, consisting of $10^{8}$ 4420 displaying cells and $10^{8}$ 3.24 displaying cells, were incubated with the 30 µL microbead-peptide mix in a 1 mL final volume for 30 minutes at room temperature with constant mixing (**Supp Fig. 2a**). Then, cells were chilled on ice for 10 minutes and subsequently washed with ice-cold PBS buffer to remove excess microbeads, biotinylated peptide and peptide-preloaded microbeads. Afterward, yeast cells were resuspended in 4 mL of ice-cold PBS buffer and run through the LS column (Miltenyi Biotec, Cat. No. 130-042-401) in the presence of a magnet. Flow through, consisting of unlabeled (not magnetized) cells, was discarded. Cells retained in the LS column were then recovered from the column in the absence of a magnet and analyzed with flow cytometry to determine their composition (**Supp Fig. 2b**). The total number of cells retained in the LS column was determined by measuring the optical density ($OD_{600}$) of eluted cells. The ratio of mCherry-expressing yeast (displaying scFv 4420) cells and EGFP-expressing yeast (displaying scFv 3.24) cells was then determined by using flow cytometry.

#### b. Screening Yeast Surface Display Libraries with Peptide-Preloaded Magnetic Beads

<u>Yeast Media and buffers used:</u>

**SD-CAA** (1 L): 5 g CAA amino acid drop-out (#DF0231172); 20 g Dextrose (#350-1); 6.7 g Yeast Nitrogen Base (# DF0919-15-3); 6.76 Citric Acid Anhydrous (# BP339-500); 11.85 Sodium citrate dihydrate (# BP327-1); pH = 4.5, filter sterilize.

**SG-CAA** (1L): 5 g CAA amino acid drop-out; 20 g Dextrose; 6.7 g Yeast Nitrogen Base; 5.4 $Na_2HPO_4$ (anhydrous) (#BP329-1); 7.46 $NaH_2PO_4$ (anhydrous) (#BP332-1); pH = 6, filter sterilize.

**YPD + NTC** (1L): 10 g Yeast Extract (#BP1422-500); 20 g Peptone (#BP14200-500); 40 g Dextrose; 100 mg NTC (#N500-500), filter sterilize.

**YPG + NTC** (1L): 10 g Yeast Extract; 20 g Peptone; 40 g Galactose (#); 100 mg NTC, filter sterilize.

**HEPES buffer** (1 L): 4.77 g HEPES (#H4034-100G); 8.76 g NaCl (#); 1 g BSA (#A3294-10G); pH = 7.4 (using 10 M NaOH), filter sterilize.

**PBS buffer** (1L):

<u>Yeast Surface Display Libraries used:</u>

Nb library v.1: diversity > $1 \times 10^{8}$; Trp selection; media used SD-CAA and SG-CAA; ref. [2]

Omni Nb library v.2: diversity > 2 $\times 10^9$; NTC selection; YPD and YPG; ref. [3]

Screening protocol used:
The following protocol describes screening yeast surface display libraries using MACS followed by FACS. Though the protocol has been optimized, additional changes could be made depending on the experimental requirements.

1. Recovering yeast libraries: Yeast surface display libraries are thawed in YPD (or any other selection media) for 2 hours at 30 °C shaker and expanded* in appropriate selective media overnight. Overnight-expanded library is then passaged** to a fresh media in order to decrease the number of dead cells. Grow passaged library overnight.

   (*: calculating the number of viable cells after thawing the library should be done with serial dilution. ** at least 10x library size should be used to passage overnight-expanded library to preserve library diversity)

2. Inducing yeast libraries: Transfer the yeast library to an induction media (YPG or SG-CAA). Yeast cells are induced at $OD_{600} = 1$ in a 20 °C water bath with shaking. We routinely induce naïve yeast libraries for 48-56 hours.

3. MACS 1st round: Prepare peptide-preloaded streptavidin Microbeads (Miltenyi Biotec #130-048-012) by mixing 100 µL beads with 10 µM biotinylated-peptide stock*. Incubate at room temperature for 1 hour with constant mixing. Peptide-preloaded microbeads are incubated** with naïve yeast display library*** for 1 hour at room temperature with constant mixing followed by incubation on ice for 15 mins. Then yeast cells are washed in order to remove excess microbeads. Washing is carried out at 4 °C using ice cold HEPES buffer. Washing is repeated twice. After the final round of washing, yeast cells are resuspended in ice cold HEPES buffer and loaded onto LS Columns (Miltenyi Biotec #130-042-401). After each loading LS Columns were washed with ice-cold HEPES buffer. After the final wash column is removed to a collection tube and cells are ejected with a plunger. This ejected yeast cells are now called Mp-I (MACS positive selection I). Mp-I cells are expanded overnight. The Mp-I library size is determined by serial dilution.

   (*bead to peptide ratio should be preserved and should be scaled accordingly. **incubation volume is such that library is entirely resuspended and final **peptide concentration** is usually 30-50 nM for the initial round. ***at least 10X library size should be used)

4. MACS 2nd round: For the second round of magnetic-activated cell sorting, 10 to 50-fold Mp-1 library size of induced cells are used. Unlike initial round, Mp-I libraries are first depleted for secondary reagent binders. Briefly, induced Mp-I library is incubated with bare magnetic beads* for 1 hour with constant mixing and chilled on ice for 15 min and subsequently washed twice with ice cold HEPES buffer. Cells were then re-suspended in ice cold HEPES buffer at $2*10^8$/mL final density and applied to pre-chilled LD column (Miltenyi, cat#130-042-901). Flow through, named Md-I (MACS depletion - I), is collected and is incubated with peptide coated Micro Beads at final concentration of 20 nM **peptide** for 1 hour at room temperature with constant mixing. Number of secondary reagent binders is calculated by eluting cells from LD column. Washing, recovery, expansion and library size calculations are carried out as in MACS round I. After the second round of magnet-activated cell sorting, library is named Mp-II (MACS positive selection – II), and recovered cells are expanded overnight. Library size is determined by serial dilution.

   (*the number of bare Micro Beads used in depletion round should be at least double the number used in positive selection round with incubation volumes at least two-fold less, essentially making the bare bead concentration 4x or more than would be in positive selection rounds, in order to efficiently remove beads binders).

5. MACS 3rd round: The third round of magnetic-activated cell sorting is carried out as in round 2 except 10 nM of final peptide concentration (coated onto Micro Beads) was used for positive selection. Library recovered after 3rd round of magnetic-activated cell sorting was named Mp-III (MACS positive selection – III). If deemed necessary, 4th and final round of magnet-activated cell sorting could also be carried out with 10 nM of final peptide concentration. Library size is determined by serial dilution as described above.

   FACS 1-5 rounds: After 3-4 rounds of magnet-activated cell sorting (MACS), we carried out fluorescence-activated cell sorting (FACS) using the BD Biosciences FACS Aria II cell sorter (University of Connecticut COR$^2$E facility). The enriched pool of yeast obtained after MACS is induced as above, and incubated with peptide pre-loaded streptavidin-PE (SAPE)* (ThermoFisher, Cat. No. S866) at room temperature with constant mixing. Peptide pre-loading is carried out at room temperature. Biotinylated peptide and SAPE are mixed at a roughly 4:1 ratio in HEPES buffer for 30-60 min with constant mixing. For SAPE,1.38 µL of SAPE stock (1 mg/mL) is mixed with 2 µL of 10 µM biotinylated-peptide stock in 10 µL of final volume. For SA-Alexa 647 (ThermoFisher, Cat. No. S32357), 0.15 µL of SA-Alexa 647 (2 mg/mL concentration) is mixed with 2 µL of 10 µM biotinylated-peptide stock in 10 µL of final volume. After pre-loading, we diluted the pre-loaded peptides to 100 - 400 nM (peptide concentration in the final solution, SA concentration would be roughly ¼) to label induced cells. During FACS, all cells with positive Nb expression and binding were collected. Between successive FACS rounds, negative selection should be carried out to remove the secondary reagent binder from the library if significant binding to secondary reagents is detected. In our case, we did not find significant secondary binding. We carried out 2-5 rounds of FACS until libraries were enriched for peptide binders.

6. Plating library and single clone analysis: After 2-5 rounds of FACS (at which point peptide binder population should be about 40-60% of library), yeast cell libraries are plated in order to obtain single clones. We picked about 100 colonies for clonal analyses. We analyzed every clone for secondary reagent as well as peptide binding. For peptide binding, we used peptide pre-loaded SAPE or/ SA-Alexa647 (peptide preloading is carried out as described above). Clones that bind to peptide preloaded SAPE or/ SA-Alexa 647 but no to SAPE or/SA-Alexa647 are then mini-preped (Zymo research, cat #D2004-A) and sequenced. For nanobodies that are going to be used for live cell imaging applications we strongly recommend expressing select clones in desired cells with fluorescence fusions before doing any characterization.

### Supplementary Figures

**a**

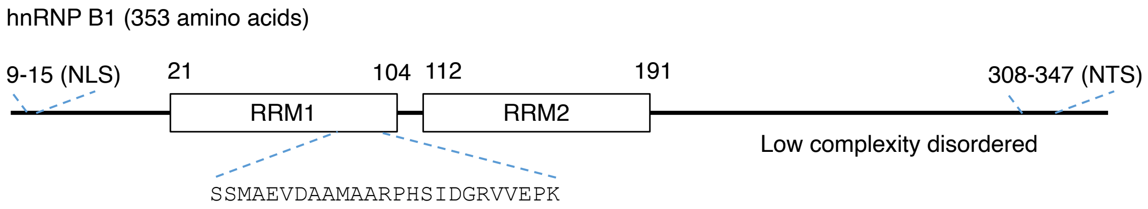


**b**

```
                                          RRM1
B1      MEKTLETVPLERKKREKEQFRKLFIGGLSFETTEESLRNYYEQWGKLTDCVVMRDPASKR 60
A2      ------------MEREKEQFRKLFIGGLSFETTEESLRNYYEQWGKLTDCVVMRDPASKR 48
                          :*****************************************
                     RRM1                                      RRM2
B1      SRGFGFVTFSSMAEVDAAMAARPHSIDGRVVEPKRAVAREESGKPGAHVTVKKLFVGGIK 120
A2      SRGFGFVTFSSMAEVDAAMAARPHSIDGRVVEPKRAVAREESGKPGAHVTVKKLFVGGIK 108
        ************************************************************
                                     RRM2
B1      EDTEEHHLRDYFEEYGKIDTIEIITDRQSGKKRGFGFVTFDDHDPVDKIVLQKYHTINGH 180
A2      EDTEEHHLRDYFEEYGKIDTIEIITDRQSGKKRGFGFVTFDDHDPVDKIVLQKYHTINGH 168
        ************************************************************
          RRM2
B1      NAEVRKALSRQEMQEVQSSRSGRGGNFGFGDSRGGGGNFGPGPGSNFRGGSDGYGSGRGF 240
A2      NAEVRKALSRQEMQEVQSSRSGRGGNFGFGDSRGGGGNFGPGPGSNFRGGSDGYGSGRGF 228
        ************************************************************

B1      GDGYNGYGGGPGGGNFGGSPGYGGGRGGYGGGGPGYGNQGGGYGGGYDNYGGGNYGSGNY 300
A2      GDGYNGYGGGPGGGNFGGSPGYGGGRGGYGGGGPGYGNQGGGYGGGYDNYGGGNYGSGNY 288
        ************************************************************

B1      NDFGNYNQQPSNYGPMKSGNFGGSRNMGGPYGGGNYGPGGSGGSGGYGGRSRY353
A2      NDFGNYNQQPSNYGPMKSGNFGGSRNMGGPYGGGNYGPGGSGGSGGYGGRSRY341
        *****************************************************
```

**Supplementary Figure 1. Peptide sequence used for screening hnRNPA2/B1 nanobody.** (**a**) Schematic of the hnRNP B1 sequence. The peptide sequence used for the nanobody screen is indicated below. NLS – nuclear localization signal (PLERKKR), NTS – nuclear targeting sequence (QQPSNYGPMKSGNFGGSRNMGGPYGGGNYGPGGSGGSGGY). (**b**) Sequence alignment of hnRNPA2 and hnRNPB1. The peptide sequence used for the nanobody screen is highlighted in yellow. RRM1 – RNA recognition motif 1, RRM2 – RNA recognition motif 2.

**a**

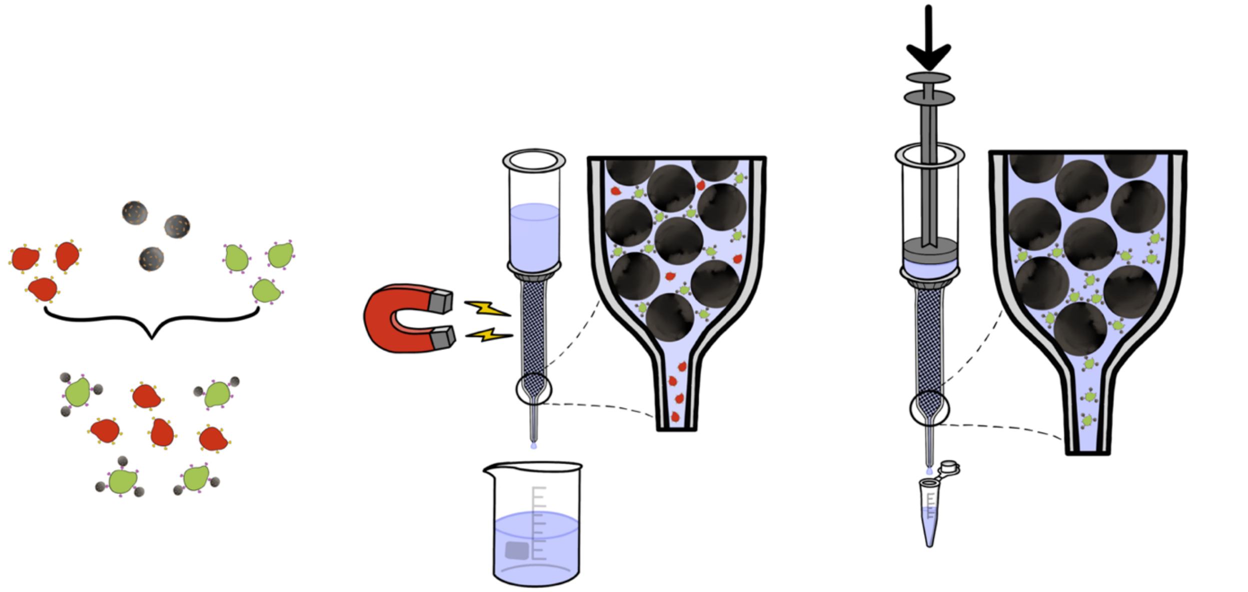

**b**

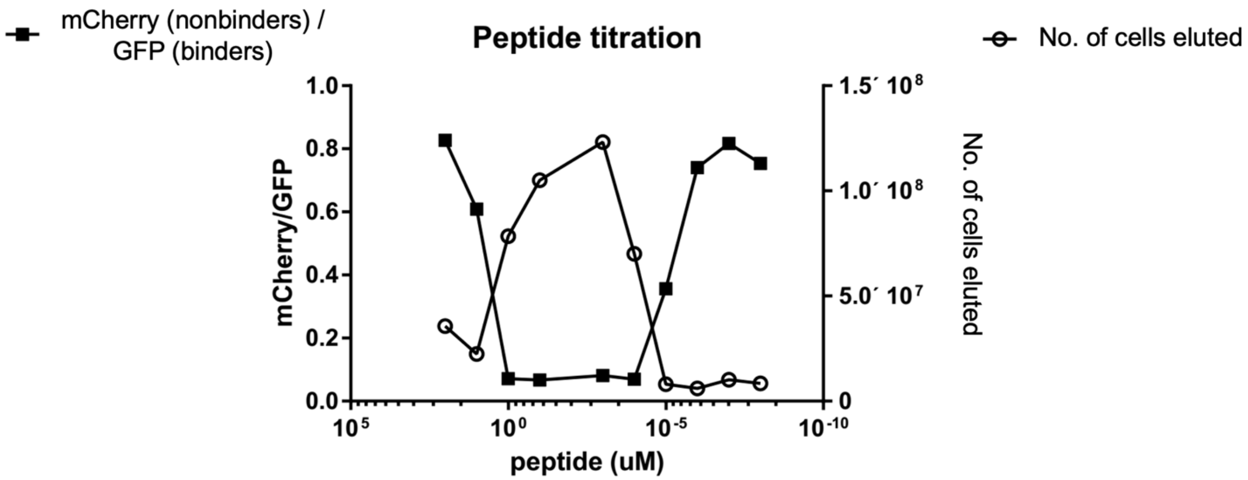


**Supplementary Figure 2. Determination of optimal peptide preloading condition. (a)** Schematic showing the overall experimental procedure. An equal number of control scFv displaying (expressed mCherry) and peptide-binding scFv 3.24 displaying cells (expressing EGFP) were incubated with peptide-magnetic microbead mix with varying concentrations of peptides. Cell mix was then loaded onto the LS column under a magnetic field, and flow-through was discarded. The remaining yeast cells in the LS column were then recovered and analyzed by flow cytometry. The total number of recovered cells was calculated by measuring optical density ($OD_{600}$). (**b**) The total number of cells eluted and the number of mCherry-expressing cells (displaying the 4420 scFv) over the number of EGFP-expressing cells (displaying the 3.24 scFv) at various peptide-microbead preloading conditions.

**Supplementary Table 1.** Nanobody sequences

| Nanobody | Sequence used in this study | Reference |
|---|---|---|
| Nb19 | MAQVQLVESGGGLVQAGGSLRLSCAASGFTFSNYAMGWYRQAPGKERELVAVINW NAGSTYADSVKGRFTISRDNAKNTVYLQMNSLKPEDTAVYYCAALAWDYVQTYDY WGQGTQVTVSS | This study |
| Nb19.3 | MARVQLVESGGGLVQAGGSLRLSCAASGFTFSSYAMGWYRQAPGKERELVAVINW NAGSTYADSVKGRFTISRDNAKNTVYLQMNSLKPEDTAAYYCAALAWDYIQTYDY WGQGTQVTVSS | This study |
| tauNb1 | MAQVQLQESGGGLVQAGGSLRLSCAASGTISYISTMGWYRQAPGKEREFVATINA GATTYYADSVKGRFTISRDNAKNTVYLQMNSLKPEDTAVYYCAVGSYAEYYLSYW GQGTQVTVSS | This study |
| cAbGFP4 | MDQVQLVESGGALVQPGGSLRLSCAASGFPVNRYSMRWYRQAPGKEREWVAGMSS AGDRSSYEDSVKGRFTISRDDARNTVYLQMNSLKPEDTAVYYCNVNVGFEYWGQG TQVTV | [4] |

Mutations in Nb19.3 are highlighted.

**Supplementary Table 2.** E3 ligase sequences and origin

| E3 ligase/ origin/brief description/ | E3 ligase protein sequence used in this study |
|---|---|
| NEDD4/Homo sapiens/aa 914-1319<br><br>GFP targeting nanobody and linker (Thr-Ser) highlighted<br><br>Check Table XYZ for cAbGFP4 Nb sequence | cAbGFP4-TS-<br>NIKRTQWEDPRLENVAITGPAVPYSRDYKRKYEFFRRKLKKQNDI<br>PNKFEMKLRRATVLEDSYRRIMGVKRADFLKARLWIEFDGEKGL<br>DYGGVAREWFFLISKEMFNPYYGLFEYSATDNYTLQINPNSGLCN<br>EDHLSYFKFIGRVAGMAVYHGKLLDGFFIRPFYKMMLHKPITLH<br>DMESVDSEYYNSLRWILENDPTELDLRFIIDEELFGQTHQHELKN<br>GGSEIVVTNKNKKEYIYLVIQWRFVNRIQKQMAAFKEGFFELIPQ<br>DLIKIFDENELELLMCGLGDVDVNDWREHTKYKNGYSANHQVIQ<br>WFWKAVLMMDSEKRIRLLQFVTGTSRVPMNGFAELYGSNGPQS<br>FTVEQWGTPEKLPRAHTCFNRLDLPPYESFEELWDKLQMAIENT<br>QGFDGVD*(stop) |
| SPOP/Homo sapiens/aa 168-374<br>GFP targeting nanobody and linker (Thr-Ser) highlighted | cAbGFP4-TS-<br>VNISGQNTMNMVKVPECRLADELGGLWENSRFTDCCLCVAGQE<br>FQAHKAILAARSPVFSAMFEHEMEESKKNRVEINDVEPEVFKEM<br>MCFIYTGKAPNLDKMADDLLAAADKYALERLKVMCEDALCSNL<br>SVENAAEILILADLHSADQLKTQAVDFINYHASDVLETSGWKSM<br>VVSHPHLVAEAYRSLASAQCPFLGPPRKRLKQS*(stop) |
| SPOP-delNLS/Homo sapiens/aa 168-369, removed C-terminal nuclear localization signal (missing aa 370-374)<br>GFP targeting nanobody and linker (Thr-Ser) highlighted | cAbGFP4-TS-<br>VNISGQNTMNMVKVPECRLADELGGLWENSRFTDCCLCVAGQE<br>FQAHKAILAARSPVFSAMFEHEMEESKKNRVEINDVEPEVFKEM<br>MCFIYTGKAPNLDKMADDLLAAADKYALERLKVMCEDALCSNL<br>SVENAAEILILADLHSADQLKTQAVDFINYHASDVLETSGWKSM<br>VVSHPHLVAEAYRSLASAQCPFLGPPR*(stop) |
| Keap1/Homo sapiens/ aa 1-324<br>GFP targeting nanobody and linker (Thr-Ser) highlighted | MQPDPRPSGAGACCRFLPLQSQCPEGAGDAVMYASTECKAEVTP<br>SQHGNRTFSYTLEDHTKQAFGIMNELRLSQQLCDVTLQVKYQDA<br>PAAQFMAHKVVLASSSPVFKAMFTNGLREQGMEVVSIEGIHPKV<br>MERLIEFAYTASISMGEKCVLHVMNGAVMYQIDSVVRACSDFLV<br>QQLDPSNAIGIANFAEQIGCVELHQRAREYIYMHFGEVAKQEEFF<br>NLSHCQLVTLISRDDLNVRCESEVFHACINWVKYDCEQRRFYVQ<br>ALLRAVRCHSLTPNFLQMQLQKCEILQSDSRCKDYLVKIFEELTL<br>HKPTQVMPCRAPKV-TS-cAbGFP4*(stop) |
| A55/Vaccinia virus/aa 1-250<br>GFP targeting nanobody and linker (Thr-Ser) highlighted | MNNSSELIAVINGFRNSGRFCDISIVINDERINAHKLILSGASEYFSI<br>LFSNNFIDSNEYEVNLSHLDYQSVNDLIDYIYGIPLSLTNDNVKYI<br>LSTADFLQIGSAITECENYILKNLCSKNCIDFYIYADKYNNKKIESA<br>SFNTILQNILRLINDENFKYLTEESMIKILSDDMLNIKNEDFAPLILI<br>KWLESTQQSCTVELLRCLRISLLSPQVIKSLYSHQLVSSIYECITFL<br>NNIAFLDESFPRYH-TS-cAbGFP4*(stop) |
| Fbxw11b/Danio rerio (Zebrafish)/ aa 1-220<br>GFP targeting nanobody and linker (Thr-Ser) highlighted | METEMEDKTLEQMNTSVMDPQTADRSPKITLIKSTFICPQVSNGP<br>LTGSRKRPSEGNYEKEKDVCIQLFDQWSEADQVEFVEHLISRMC<br>HYQHGHINSYLKPMLQRDFITALPAQGLDHIAENILSFLDARSLCS<br>AELVCKEWQRVISEGMLWKKLIERMVRTDPLWKGLSERHQWEK<br>YLFKNRTTEVPPNSYYRSLYPKIIQDIETIEANWRCGRHMDQ-TS-<br>cAbGFP4*(stop) |
| BTRC (Fbxw1A)/Homo sapiens/ aa 1-286 | MDPAEAVLQEKALKFMCSMPRSLWLGCSSLADSMPSLRCLYNP<br>GTGALTAFQNSSEREDCNNGEPPRKIIPEKNSLRQTYNSCARLCLN<br>QETVCLASTAMKTENCVAKTKLANGTSSMIVPKQRKLSASYEKE<br>KELCVKYFEQWSESDQVEFVEHLISQMCHYQHGHINSYLKPMLQ<br>RDFITALPARGLDHIAENILSYLDAKSLCAAELVCKEWYRVTSDG |

| | |
|---|---|
| GFP targeting nanobody and linker (Thr-Ser) highlighted | MLWKKLIERMVRTDSLWRGLAERRGWGQYLFKNKPPDGNAPPN SFYRALYPKIIQDIETIESNW-TS-cAbGFP4*(stop) |
| Fbxw 11 Isform B/Homo sapiens/ aa 1-218<br>GFP targeting nanobody and linker (Thr-Ser) highlighted | MEPDSVIEDKTIELMNTSVMEDQNEDESPKKNTLWQISNGTSSVI VSRKRPSEGNYQKEKDLCIKYFDQWSESDQVEFVEHLISRMCHY QHGHINSYLKPMLQRDFITALPEQGLDHIAENILSYLDARSLCAAE LVCKEWQRVISEGMLWKKLIERMVRTDPLWKGLSERRGWDQYL FKNRPTDGPPNSFYRSLYPKIIQDIETIESNWRCGRHNLQ-TS-cAbGFP4*(stop) |
| Fbxw 7 Isoform A/Homo sapiens/ aa 1-367<br>GFP targeting nanobody and linker (Thr-Ser) highlighted | MNQELLSVGSKRRRTGGSLRGNPSSSQVDEEQMNRVVEEEQQQ QLRQQEEEHTARNGEVVGVEPRPGGQNDSQQGQLEENNNRFISV DEDSSGNQEEQEEDEEHAGEQDEEDEEEEEMDQESDDFDQSDDS SREDEHTHTNSVTNSSSIVDLPVHQLSSPFYTKTTKMKRKLDHGS EVRSFSLGKKPCKVSEYTSTTGLVPCSATPTTFGDLRAANGQGQQ RRRITSVQPPTGLQEWLKMFQSWSGPEKLLALDELIDSCEPTQVK HMMQVIEPQFQRDFISLLPKELALYVLSFLEPKDLLQAAQTCRYW RILAEDNLLWREKCKEEGIDEPLHIKRRKVIKPGFIHSPWKSAYIR QHRIDTNWRR-TS-cAbGFP4*(stop) |
| IpaH9.8/Shigella flexneri/ aa 254-545<br>GFP targeting nanobody and linker (Thr-Ser) highlighted | cAbGFP4-TS-LADAVTAWFPENKQSDVSQIWHAFEHEEHANTFSAFLDRLSDTV SARNTSGFREQVAAWLEKLSASAELRQQSFAVAADATESCEDRV ALTWNNLRKTLLVHQASEGLFDNDTGALLSLGREMFRLEILEDIA RDKVRTLHFVDEIEVYLAFQTMLAEKLQLSTAVKEMRFYGVSGV TANDLRTAEAMVRSREENEFTDWFSLWGPWHAVLKRTEADRW AQAEEQKYEMLENEYPQRVADRLKASGLSGDADAEREAGAQV MRETEQQIYRQLTDEVLALRLPENGSQLHHS*(stop) |
| TRIM21/Mus musculus/ aa 9-470<br><br>GFP targeting nanobody and linker (Thr-Ser) highlighted | MSLEKMWEEVTCSICLDPMVEPMSIECGHCFCKECIFEVGKNGGS SCPECRQQFLLRNLRPNRHIANMVENLKQIAQNTKKSTQETHCM KHGEKLHLFCEEDGQALCWVCAQSGKHRDHTRVPIEEAAKVYQ EKIHVALEKLRKGKELAEKMEMDLTMQRTDWKRNIDTQKSRIH AEFALQNSLLAQEEQRQLQRLEKDQREYLRLLGKKEAELAEKNQ ALQELISELERRIRGSELELLQEVRIILERSGSWNLDTLDIDAPDLT STCPVPGRKKMLRTCWVHITLDRNTANSWLIISKDRRQVRMGDT HQNVSDNKERFSNYPMVLGAQRFSSGKMYWEVDVTQKEAWDL GVCRDSVQRKGQFSLSPENGFWTIWLWQDSYEAGTSPQTTLHIQ VPPCQIGIFVDYEAGVVSFYNITDHGSLIYTFSECVFAGPLRPFFNV GFNYSGGNAAPLKLCPLKM-TS-cAbGFP4*(stop) |
| TRIM21/Mus musculus/ aa 9-470 with B-Box like deletion (missing aa 95-126)<br>GFP targeting nanobody and linker (Thr-Ser) highlighted | MSLEKMWEEVTCSICLDPMVEPMSIECGHCFCKECIFEVGKNGGS SCPECRQQFLLRNLRPNRHIANMVENLKQIAQNTKKSTQETHTRV PIEEAAKVYQEKIHVALEKLRKGKELAEKMEMDLTMQRTDWKR NIDTQKSRIHAEFALQNSLLAQEEQRQLQRLEKDQREYLRLLGKK EAELAEKNQALQELISELERRIRGSELELLQEVRIILERSGSWNLDT LDIDAPDLTSTCPVPGRKKMLRTCWVHITLDRNTANSWLIISKDR RQVRMGDTHQNVSDNKERFSNYPMVLGAQRFSSGKMYWEVDV TQKEAWDLGVCRDSVQRKGQFSLSPENGFWTIWLWQDSYEAGT SPQTTLHIQVPPCQIGIFVDYEAGVVSFYNITDHGSLIYTFSECVFA GPLRPFFNVGFNYSGGNAAPLKLCPLKM-TS-cAbGFP4*(stop) |
| TRIM21/Mus musculus/ aa 9-271 | MSLEKMWEEVTCSICLDPMVEPMSIECGHCFCKECIFEVGKNGGS SCPECRQQFLLRNLRPNRHIANMVENLKQIAQNTKKSTQETHCM KHGEKLHLFCEEDGQALCWVCAQSGKHRDHTRVPIEEAAKVYQ EKIHVALEKLRKGKELAEKMEMDLTMQRTDWKRNIDTQKSRIH AEFALQNSLLAQEEQRQLQRLEKDQREYLRLLGKKEAELAEKNQ ALQELISELERRIRGSELELLQEVRIILERSGSWNLDTLDIDAP-TS-cAbGFP4*(stop) |

| | |
|---|---|
| TRIM21/Mus musculus/ aa 9-271 with B-Box like deletion (missing aa 95-126) | MSLEKMWEEVTCSICLDPMVEPMSIECGHCFCKECIFEVGKNGGS SCPECRQQFLLRNLRPNRHIANMVENLKQIAQNTKKSTQETHTRV PIEEAAKVYQEKIHVALEKLRKGKELAEKMEMDLTMQRTDWKR NIDTQKSRIHAEFALQNSLLAQEEQRQLQRLEKDQREYLRLLGKK EAELAEKNQALQELISELERRIRGSELELLQEVRIILERSGSWNLDT LDIDAP-TS-cAbGFP4*(stop) |

**Supplementary Table 3.** Nb19 clones with unique mutations identified from isolated clones.

| **NbR3S2 clone** | **Mutations (CDR mutations in bold)** |
|---|---|
| Nb19.1 | **N32S**; T78A; **V104I**; **Q105R** |
| Nb19.3 | Q2R; **N32S**; V93A; **V104I** |
| Nb19.4 | Q2R; **N32S**; **V104I** |
| Nb19.6 | **S31P**; **N32S**; **T59A**; I70V; N84S; **V104I** |
| Nb19.7 | **N32S**; **N53D**; **V104I** |
| Nb19.10 | A25T; **S31P**; I70V; **V104**I |
| Nb19.11 | **N32S**; **N55D**; A61V; **V104I** |
| Nb19.12 | Q2R; **N32S**; **I52T**; D62N; **V104I** |
| Nb19.13 | Q2R; **N32S**; **S58P**; T78A; **V104I** |
| Nb19.14 | **N32S**; N74D; T78A; **V104I** |
| Nb19.16 | V6A; **N32S**; **V104I** |
| Nb19.17 | **N32S**; Y95D; **V104I** |
| Nb19.19 | R20H; A25T; **N32S**; A75V; M83T; **V104I** |
| Nb19.20 | L21S; **N32S**; **V51A**; **N53D**; K87E; E89G; **V104I** |
| Nb19.21 | **N32S**; **V51I**; K65E; N84D; V93A; **V104I** |
| Nb19.22 | **S31P**; **N32S**; **V51A**; **V104I**; V116A |
| Nb19.23 | Q2R; **N32S**; E45G; **N55S**; **V104I** |
| Nb19.24 | **N32S**; **G57S**; K87E; E89G; **V104I** |
| Nb19.28 | S18N; **N32S**; **V104I** |
| Nb19.30 | **N32S**; **V104I** |

### **Supplementary Table 4.** Primer sequences

| **Primer or Plasmid Name /Brief description** | **Sequence** | **Reference/Addgene #/** |
|---|---|---|
| pYDS649 v.1 and pYDS649 v.2 sequencing primers | Forward: TAATATACCTCTATACTTTAACGTCAAGG<br>Reverse: TACTGATGCTTCTGTAGAGGGTGAGG | [2], [3] |
| pCMV sequencing primers | Forward: AAGCTTCGAATTCTGC<br>Reverse: CATTTTATGTTTCAGGTTCAGGG | This work |
| pCMV-cAbGFP-mCherry | Forward-spacer-BamHI-Kozak-cAbGFP overlap:<br>AAAT-GGATCC-GCCACC-ATGGATCAAGTCCAACTGGT<br>Reverse-spacer-SpeI-cAbGFP overlap:<br>AAAT-ACTAGT-GACGGTGACCTG<br>Forward-spacer-SpeI-linker-mCherry overlap:<br>AAAT-ACTAGT-TCCAGCGGCTCTGGTGGC-ATGGTTTCAAAGGGCGAAG<br>Reverse-Spacer-BsrGI-Stop-mCherry overlap:<br>AAAT-TGTACA-TCA-CTTATACAATTCATCCATACC | This work<br><br>(cAbGFP was obtain from [4]) |
| pCMV-EGFP-hnRNPA2 (WT)<br><br>Expresses EGFP-hnRNPA2 (wild type) | Reverse-spacer- BsaI recognition and cut site-backbone overlap:<br>AAAT-GGTCTC T GATC-GATCCCGGGCCCGC<br>Forward-spacer-BsaI recognition and cut site-Kozak-EGFP overlap:<br>AAAT-GGTCTC T GATC-CGCCACC-ATGGTGAGCAAGGGCGAG<br>Reverse-spacer- BsaI recognition and cut site-EGFP overlap:<br>AAAT-GGTCTC T CTTG-TACAGCTCGTCCATG<br>Forward-spacer- BsaI recognition and cut site-hnRNPA2 overlap:<br>AAAT-GGTCTC T CAAG-ATGGAACGTGAAAAAGAGCAG<br>Reverse-spacer- BsaI recognition and cut site-stop-hnRNPA2 overlap:<br>AAAT-GGTCTC T CCGC-TCAGTAACGGCTGCGGCCAC<br>Forward-spacer- BsaI recognition and cut site-backbone overlap:<br>AAAT-GGTCTC T GCGG-CCGCGACTCTAG | This work<br><br><br>(hnRNPA2 was obtain from Addgene #139109) |
| pCMV-NES$^{PKI}$-EGFP-mCherry<br><br>Made from:pCMV-cAbGFP-mCherry by swapping cAbGFP4 with NES-EGFP<br><br>Description: EGFP has nuclear export signal (NES) from protein kinase A inhibitor. | Forward-Spacer-BamHI-Kozak-NES$^{PKI}$-EGFP overlap:<br>AAAT- GGATCC-GCCACC-ATGAACTCCAACGAACTCGCTCTGAAGCTTGCTGGTCTGGACATCAACAAG-ATGGTGAGCAAGGGCGAG<br>Reverse-spacer-SpeI-EGFP overlap:<br>AAAT-ACTAGT-CTTGTACAGCTCGTCCATG<br><br>NES$^{PKI}$ amino acid sequence: NSNELALKLAGLDINK | This work |
| pCMV-NES$^{PKI}$-EGFP-hnRNPA2 (WT)<br><br>Made from: pCMV-EGFP-hnRNPA2 (WT).<br><br>Description: EGFP has nuclear export signal (NES) from protein kinase A inhibitor. | Forward-Spacer-BamHI-Kozak-NES$^{PKI}$-EGFP overlap:<br>AAAT- GGATCC-GCCACC-ATGAACTCCAACGAACTCGCTCTGAAGCTTGCTGGTCTGGACATCAACAAG-ATGGTGAGCAAGGGCGAG<br>Reverse-spacer-NotI-stop-hnRNPA overlap:<br>AAAT-CGCGGCCGC-TCA-GTAACGGCTGCGGCCAC | This work |
| pCMV-NES$^{PKI}$-EGFP-EGFP-hnRNPA2 (WT)<br><br>Made from: NES$^{PKI}$-EGFP-mCherry by swapping mCherry with EGFP-hnRNPA2B1<br><br>Description: Exports two EGFP fused to hnRNPA2 (WT) | Forward-Spacer-SpeI-linker-EGFP overlap:<br>AAAT-ACTAGT-TCCAGCGGCTCTGGTGGC-ATGGTGAGCAAGGGCGAGGAGCTGTT<br><br>Reverse-spacer-NotI-stop-hnRNPA overlap:<br>AAAT-CGCGGCCGC-TCA-GTAACGGCTGCGGCCAC | This work |

| | | |
|---|---|---|
| pCMV-NES$^{PKI}$-EGFP-hnRNPA2 delLCD<br><br>Made from: pCMV-NES$^{PKI}$-EGFP-hnRNPA2 (WT)<br><br>Description: Expresses hnRNPA2 without Low Complexity Domain (delLCD). | Forward-Spacer-BamHI-Kozak-NES$^{PKI}$-EGFP overlap:<br>AAAT- GGATCC-GCCACC-<br>ATGAACTCCAACGAACTCGCTCTGAAGCTTGCTGGTCTGGACATCA<br>ACAAG-ATGGTGAGCAAGGGCGAG<br><br>Reverse-spacer-NotI-stop-hnRNPA overlap:<br>AAAT-GCGGCCGC-TCA-CTCTTGACGGCTCAGCG | This work |
| pCMV-NES$^{PKI}$-EGFP-EGFP-hnRNPA2 delLCD<br><br>Made from: NES$^{PKI}$-EGFP-mCherry by swapping mCherry with EGFP-hnRNPA2 delLCD<br><br>Description: Exports two EGFP fused to hnRNPA2 delLCD | Forward-Spacer-SpeI-linker-EGFP overlap:<br>AAAT-ACTAGT-TCCAGCGGCTCTGGTGGC-<br>ATGGTGAGCAAGGGCGAGGAGCTGTT<br><br>Reverse-spacer-NotI-stop-hnRNPA overlap:<br>AAAT-GCGGCCGC-TCA-CTCTTGACGGCTCAGCG | This work |
| pCMV-NES$^{PKI}$-EGFP-hnRNPA2 del PY-NLS<br><br>Made from: pCMV-NES$^{PKI}$-EGFP-hnRNPA2 (WT)<br><br>Description: Expresses EGFP-hnRNPA2 without PY-NLS | Forward-Spacer-BamHI-Kozak-NES$^{PKI}$-EGFP overlap:<br>AAAT- GGATCC-GCCACC-<br>ATGAACTCCAACGAACTCGCTCTGAAGCTTGCTGGTCTGGACATCA<br>ACAAG-ATGGTGAGCAAGGGCGAG<br><br>Reverse-spacer-NotI-stop-hnRNPA overlap:<br>AAAT-GCGGCCGC-TCA-GTTGTAGTTGCCAAAATCATTG | This work |
| pCMV-NES$^{PKI}$-EGFP-EGFP-hnRNPA2 del PY-NLS<br><br>Made from: NES$^{PKI}$-EGFP-mCherry by swapping mCherry with EGFP-hnRNPA2 del PY-NLS<br><br>Description: Expresses EGFP-EGFP-hnRNPA2 without PY-NLS | Forward-Spacer-SpeI-linker-EGFP overlap:<br>AAAT-ACTAGT-TCCAGCGGCTCTGGTGGC-<br>ATGGTGAGCAAGGGCGAGGAGCTGTT<br>Reverse-spacer-NotI-stop-hnRNPA overlap:<br>AAAT-GCGGCCGC-TCA-GTTGTAGTTGCCAAAATCATTG | |
| pCMV-Nb#19-mCherry<br><br>Made from: pCMV-cAbGFP-mCherry<br>Descriotion: Expresses hnRNPA2B1 nanobody clone #19 with mCherry fusion | Forward-spacer-BamHI-Kozak-Nb 19 overlap:<br>AAAT-GGATCC-GCCACC-ATGGCACAAGTTCAGCTTGTAGAG<br>Reverse-spacer-SpeI-cAbGFP overlap:<br>AAAT-ACTAGT-CGATGATACAGTTACTTGGG | This work Nanobody #19 was PCR amplified from pYDS649 v.2 |
| pCMV-di(Nb#19)-mCherry<br><br>Made from: pCMV-cAbGFP-mCherry<br>Descriotion: Expresses tandem dimeric hnRNPA2B1 nanobody clone #19 with mCherry fusion | Reverse-spacer- BsaI recognition and cut site-backbone overlap:<br>AAAT-GGTCTC T GGTG-GCGGATCCCGG<br>Forward-spacer- BsaI recognition and cut site-1$^{st}$ Nb #19 overlap:<br>AAAT-GGTCTC T CACC-ATGGCACAAGTTCAGCTTGTAGAG<br>Reverse-spacer- BsaI recognition and cut site-SG linker -1$^{st}$ Nb #19 overlap<br>AAAT-GGTCTC T TCCG-<br>GAACCACCACCGCCGCTGCCACCGCCACCGGA-<br>CGATGATACAGTTACTTGGGTAC<br>Forward-spacer- BsaI recognition and cut site-SG linker-2$^{nd}$ Nb #19 overlap:<br>AAAT-GGTCTC T CGGA-<br>GGCGGCGGTAGCGGCGGAGGTGGAGCACAAGTTCAGCTTGTAGAG<br>Reverse-spacer- BsaI recognition and cut site-2$^{nd}$ Nb #19 overlap:<br>AAAT-GGTCTC T TAGT-CGATGATACAGTTACTTGGG<br>Forward-spacer- BsaI recognition and cut site-SG linker mCherry overlap:<br>AAAT-GGTCTC T ACTA-<br>GTTCCAGCGGCTCTGGTGGCATGGTTTCAAAGGGCGAAG | This work |
| pCMV-tri(Nb#19)-mCherry | Forward-spacer- BsaI recognition and cut site-1$^{st}$ Nb #19 overlap:<br>AAAT-GGTCTC T CACC-ATGGCACAAGTTCAGCTTGTAGAG | This work |

| | | |
|---|---|---|
| Made from: pCMV-cAbGFP-mCherry<br>Descriotion: Expresses tandem trimeric hnRNPA2B1 nanobody clone #19 with mCherry fusion | Reverse-spacer- BsaI recognition and cut site-SG linker -1st Nb #19 overlap<br>AAAT-GGTCTC T TCCG-GAACCACCACCGCCGCTGCCACCGCCACCGGA-CGATGATACAGTTACTTGGGTAC<br>Forward-spacer- BsaI recognition and cut site-SG linker-2nd Nb #19 overlap:<br>AAAT-GGTCTC T CGGA-GGCGGCGGTAGCGGCGGAGGTGGAGCACAAGTTCAGCTTGTAGAG<br>Reverse-spacer- BsaI recognition and cut site-SG linker-2nd Nb #19 overlap:<br>AAAT-GGTCTC T CGGA-ACCACCACCGCCGCTGCCACCGCCACCGGACGATGATACAGTTACTTGGGTAC<br>Forward-spacer- BsaI recognition and cut site-SG linker-3rd Nb #19 overlap:<br>AAAT-GGTCTC T CGGA-TCCGGAGGCGGCGGTAGCGGCGGAGGTGGAGCACAAGTTCAGCTTGTAGAG<br>Reverse-spacer- BsaI recognition and cut site-3rd Nb #19 overlap:<br>AAAT-GGTCTC T TAGT-CGATGATACAGTTACTTGGG<br>Forward-spacer- BsaI recognition and cut site-SG linker mCherry overlap:<br>AAAT-GGTCTC T ACTA-GTTCCAGCGGCTCTGGTGGCATGGTTTCAAAGGGCGAAG | |
| pCMV-cAbGFP-NEDD4<br><br>Made from: pCMV-cAbGFP-mCherry<br>Description: Express murine NEDD4 targeting EGFP (amino acid sequence given in SUPP. Table XYZ) | Forward-spacer-SpeI-NEDD 4 overlap:<br>AAAT-ACTAGT-AATATAAAAAGAACACAATGGGAAGATC<br>Reverse-spacer-BsrGI-stop-NEDD 4 overlap:<br>AAAT-TGTACA-CTA-ATCAACTCCATCAAAGCCCTG | mNEDD4 from Addgene #38316<br><br>This work |
| pCMV-cAbGFP-SPOP<br><br>Made from: pCMV-cAbGFP-mCherry<br>Description: Express SPOP targeting EGFP (amino acid sequence given in SUPP. Table XYZ) | Forward-spacer-SpeI-SPOP overlap:<br>AAAT-ACTAGT-GTCAACATTTCTGGCCAGAAT<br>Reverse-spacer-BsrGI-stop-SPOP overlap:<br>AAAT-TGTACA-TTA-GGATTGCTTCAGGCGTTT | SPOP coding g-block was synthesized with Integrated DNA Technologies Inc.<br><br>This work |
| pCMV-cAbGFP-SPOPdelNLS<br><br>Made from: pCMV-cAbGFP-mCherry<br>Description: Express SPOP targeting EGFP (amino acid sequence given in SUPP. Table XYZ) | Forward-spacer-SpeI-SPOP overlap:<br>AAAT-ACTAGT-GTCAACATTTCTGGCCAGAAT<br><br>Reverse-spacer-BsrGI-stop-SPOP overlap:<br>AAAT-TGTACA-TTA-GCGTGGGGGTCCCAG | This work |
| pCMV-Keap1-cAbGFP4<br><br>Description: Express Keap1 targeting EGFP (amino acid sequence given in SUPP. Table XYZ) | Forward-spacer-BamHI-Kozak-Keap1 overlap:<br>AAAT-GGATCC-GCCACC-ATGCAGCCAGATCCCAGGCCTAGCGGGGCTGGG<br>Reverse-spacer-SpeI-Keap 1 overlap<br>AAAT-ACTAGT-CACCTTGGGCGCCCG<br><br>Forward-spacer-SpeI-cAbGFP overlap:<br>AAAT-ACTAGT-ATGGATCAAGTCCAACTGGTG<br>Reverse-spacer-BsrGI-stop-cAbGFP4 overlap:<br>AAAT-TGTACA-CTA-AGTGACGGTGACCTGGGTG | Keap 1 coding g-block was synthesized with Integrated DNA Technologies Inc.<br><br>This work |
| pCMV-A55-cAbGFP<br><br>Description: Express A55 targeting EGFP (amino acid sequence given in SUPP. Table XYZ) | Forward-spacer-SpeI-A55 overlap:<br>AAAT-GGATCC-GCCACC-GGATCCGCCACCATGAACAACTCCTCCGAGCT<br>Reverse-spacer-SpeI-A55 overlap<br>AAAT-ACTAGT-GTGGTAGCGGGGGAAGG | A55 coding g-block was synthesized with Integrated DNA Technologies Inc.<br><br>This work |
| pCMV-Fbxw11b-cAbGFP<br>Description: Express Fbxw11b targeting EGFP (amino acid sequence given in SUPP. Table XYZ) | Forward-spacer-BamHI-Kozak- Fbxw11b overlap:<br>AAAT-GGATCC-GCCACC-ATGGAGACGGAGATGGAGG<br>Reverse-spacer-SpeI-Fbxw11b overlap<br>AAAT-ACTAGT-TTGATCCATATGTCGACCACAC<br>Forward-spacer-SpeI-cAbGFP overlap: | Fbxw11b from Addgene #119716<br><br>This work |

| | | |
|---|---|---|
| | AAAT-ACTAGT-ATGGATCAAGTCCAACTGGTG<br>Reverse-spacer-BsrGI-stop-cAbGFP4 overlap:<br>AAAT-TGTACA-CTA-AGTGACGGTGACCTGGGTG | |
| pCMV-Fbxw1-cAbGFP<br><br>Description: Express Fbxw1 targeting EGFP (amino acid sequence given in SUPP. Table XYZ) | Forward-spacer-BamHI-Kozak- Fbxw1 overlap:<br>AAAT-GGATCC-GCCACC-ATGGACCCGGCCGAGG<br>Reverse-spacer-SpeI-Fbxw1 overlap:<br>AAAT-ACTAGTCCAATTAGATTCTATTGTCTCAATGTC<br>Forward-spacer-SpeI-cAbGFP overlap:<br>AAAT-ACTAGT-ATGGATCAAGTCCAACTGGTG<br>Reverse-spacer-BsrGI-stop-cAbGFP4 overlap:<br>AAAT-TGTACA-CTA-AGTGACGGTGACCTGGGTG | Fbxw1 from Addgene #82244<br><br>This work |
| pCMV-Fbxw11-cAbGFP<br><br>Description: Express Fbxw11 targeting EGFP (amino acid sequence given in SUPP. Table XYZ) | Forward-spacer-BamHI-Kozak- Fbxw11 overlap:<br>AAAT-GGATCC-GCCACC-ATGGAGCCAGATAGCGTTATAG<br>Reverse-spacer-SpeI-Fbxw11 overlap:<br>AAAT-ACTAGT-CTGCAGGTTATGGCGCC<br>Forward-spacer-SpeI-cAbGFP overlap:<br>AAAT-ACTAGT-ATGGATCAAGTCCAACTGGTG<br>Reverse-spacer-BsrGI-stop-cAbGFP4 overlap:<br>AAAT-TGTACA-CTA-AGTGACGGTGACCTGGGTG | Fbxw11 coding g-block was synthesized with Integrated DNA Technologies Inc.<br><br>This work |
| pCMV-Fbxw7-cAbGFP4<br><br>Description: Express Fbxw7 targeting EGFP (amino acid sequence given in SUPP. Table XYZ) | Forward-spacer-BamHI-Kozak- Fbxw7 overlap:<br>AAAT-GGATCCGCCACCATGAATCAGGAACTGCTCTCTGTG<br>Reverse-spacer-SpeI-Fbxw11 overlap:<br>AAAT-ACTAGT-TCGCCTCCAGTTAGTATCAATTC<br>Forward-spacer-SpeI-cAbGFP overlap:<br>AAAT-ACTAGT-ATGGATCAAGTCCAACTGGTG<br>Reverse-spacer-BsrGI-stop-cAbGFP4 overlap:<br>AAAT-TGTACA-CTA-AGTGACGGTGACCTGGGTG | Fbxw7 from Addgene #81795<br><br>This work |
| pCMV-cAbGFP-IpaH9.8<br><br>Made from: pCMV-cAbGFP-mCherry by swapping mCherry with IpaH9.8<br>Description: Express IpaH9.8 targeting EGFP (amino acid sequence given in SUPP. Table XYZ) | Forward-spacer-SpeI-IpaH9.8 overlap:<br>AAAT-ACTAGT-TTAGCCGATGCCGTGAC<br>Reverse-spacer-BsrGI-stop-IpaH9.8 overlap:<br>AAAT-TGTACA-CTA-GCTGTGGTGAAGTTGTGATCC | IpaH9.8 coding g-block was synthesized with Integrated DNA Technologies Inc.<br><br>This work |
| pCMV-mTRIM21-cAbGFP4<br>Description: Express full length murine TRIM21 targeting EGFP (amino acid sequence given in SUPP. Table XYZ) | Forward-spacer-EcorI-Kozak-TRIM21 overlap:<br>AAAT-GAATTC-AGCCACC-ATGTCTCTGGAAAAGATGTGG<br>Reverse-spacer-SpeI-TRIM21 overlap:<br>AAAT-ACTAGT-CATCTTTAGTGGACAGAGCTTTAG | murine TRIM21 from Addgene #105516<br><br>This work |
| pCMV-mTRIM21delBBox-cAbGFP<br><br>Made from: pCMV-mTRIM21-cAbGFP4<br><br>Description: Express murine TRIM21 without B-Box like domain (amino acid sequence given in SUPP. Table XYZ) | Reverse-BsaI recognition site-TRIM21 overlap:<br>AAAT-GGGTCTC T GTGC-GTTTCCTGGGTACTC<br>Forward-BsaI recognition site-TRIM21 overlap:<br>AAAT-GGTCTC T GCAC-ACCAGGGTCCCTATTGAAG | This work |
| pCMV-mTRIM21delB30.2/SPRY-cAbGFP4<br><br>Made from: pCMV-mTRIM21-cAbGFP4<br><br>Description: Express murine TRIM21 without B30.2/SPRY (amino acid sequence given in SUPP. Table XYZ) | Forward-spacer-EcorI-Kozak-TRIM21 overlap:<br>AAAT-GAATTC-AGCCACC-ATGTCTCTGGAAAAGATGTGG<br>Reverse-spacer-SpeI-TRIM21 overlap:<br>AAAT-ACTAGT-TGGGGCGTCAATATCTAACGTG | This work |

| | | |
|---|---|---|
| pCMV-mTRIM21delBBox-delB30.2/SPRY-cAbGFP4<br><br>Made from: pCMV-mTRIM21delB30.2/SPRY-cAbGFP4<br><br>Description: Express murine TRIM21 without B30.2/SPRY (amino acid sequence given in SUPP. Table XYZ) | Reverse-BsaI recognition site-TRIM21 overlap:<br>AAAT-GGGTCTC T GTGC-GTTTCCTGGGTACTC<br>Forward-BsaI recognition site-TRIM21 overlap:<br>AAAT-GGTCTC T GCAC-ACCAGGGTCCCTATTGAAG | This work |
| pCMV-mLnx1-cAbGFP4<br><br>Description: Express murine Lnx1 (amino acid sequence given in SUPP. Table XYZ) | Forward-spacer-BamHI-Kozak-Lnx1 overlap:<br>AAAT-GGATCC-GCCACC-ATGAATCAGCCTGACCTTG<br>Reverse-spacer-SpeI-Lnx1 overlap:<br>AAAT-ACTAGT-GGCAGCGGCACTAACTTC | murine Lnx1 coding g-block was synthesized with Integrated DNA Technologies Inc.<br><br>This work |
| F_homology2 | ATCTTCGCTGCTTTGCCATTGGCCTTGGCTGATTACAAGGACGATGACGATAAGGCTAGC | This work |
| R_homology2 | ACCTCTACCTTCAATGGCGGCCGCCAAGTCTTCTTCAGAAATAAGCTTTTGTTCTGTACA | This work |
